# SteerAF: Distogram-based Steering of AlphaFold2 toward Alternative Conformations

**DOI:** 10.64898/2026.06.19.733296

**Authors:** Jiajun Tang, Zefeng Zhu, Song Yang, Chen Song

## Abstract

End-to-end structure predictors, such as AlphaFold2, typically output only the dominant conformational state of a given protein, which is biased by the training dataset. Existing strategies for recovering alternative conformations are often computationally expensive and offer limited biological interpretability. Here, we present SteerAF, an inference-time optimization framework based on AlphaFold2 that leverages information encoded in the distogram derived from deep multiple sequence alignments (MSAs) to predict alternative protein conformations. Across four benchmark datasets, SteerAF matches or surpasses existing methods in predicting alternative conformations for the majority of systems. Sparse MSA-feature modifications generated via block gradient ascent exhibit a strong correlation with experimentally characterized functional residues, recovering them with approximately 50% precision in the tested proteins. Furthermore, SteerAF enables effective decoy selection in the absence of experimental structures, and its predictions can serve as seed structures for molecular dynamics simulations to map conformational landscapes. Thus, SteerAF provides an efficient and interpretable approach for predicting alternative conformations, offering a framework that can be extended to other similar predictors and problems.

## 1 Introduction

The remarkable success of end-to-end protein structure prediction (PSP) models such as AlphaFold2^1^ has been enabled by two converging resources: the abundant co-evolutionary information embedded in rapidly expanding sequence databases, and the wealth of experimentally determined structures accumulated in the Protein Data Bank. ^2–4^ From this perspective, AlphaFold2 essentially solves the problem of mapping the co-evolutionary restraints encoded in a multiple sequence alignment (MSA) onto a precise three-dimensional structure. However, biological reality is far more complex than a single static conformation. Proteins are inherently dynamic macromolecules that exist as conformational ensembles governed by the Boltzmann distribution. Crucial processes such as enzymatic catalysis, receptor signal transduction, antibody–antigen recognition, and allosteric regulation fundamentally rely on structural transitions among multiple conformations. However, regression-based models (e.g. AlphaFold2 and RoseTTAFold ^5^) predominantly predict the single dominant conformation most frequently observed in the training data. ^6^ Consequently, developing robust methods for multi-conformation prediction has become a pivotal challenge in structural biology.

Current multi-conformation prediction strategies fall into three main categories. (i) MSA prompting: techniques that alter MSA inputs to generate alternative conformations, e.g., by subsampling the alignment, ^7–9^ stochastically masking MSA inputs, ^10,11^ or clustering MSA entries ^12,13^ to bias the outputs. (ii) Internal representation manipulation: by modulating intermediate hidden states with or without auxiliary information ^14–16^ to bias the outputs. (iii) Generative modeling: approaches that explicitly learn the conditional probability distribution *p*(structure | sequence) to sample conformational ensembles approximating the Boltzmann distribution, ^17–20^ or that bias the diffusion process to guide exploration toward specific conformational states via enhanced sampling. ^21^ However, most of these approaches suffer from intensive sampling and require substantial computational time. Molecular dynamics (MD) simulations also contribute to this field, but pure MD-based methods are even more computationally expensive. Consequently, they are increasingly supplanted by, or integrated with, the above prediction strategies, typically deployed only when essential for validation or refinement. ^22–24^ Experimental restraints can also be incorporated into the prediction pipelines for predicting alternative conformations. ^25–31^ However, system-tailored experimental restraints are not always readily available.

In light of early studies of direct coupling analysis, ^32,33^ our prior works ^34,35^ examined per-residue-pair contact and distance-bin probability distributions (distograms) from PSP models. By applying empirical rules to filter these signals, combined with Rosetta-based modeling, we reconstructed alternative conformations for most targets in two-state conformational benchmark datasets. These results demonstrated that predicted contact maps and distograms derived from deep MSAs implicitly encode signatures of alternative conformational states, opening a route for predicting alternative conformations in a self-supervised manner.

In this study, we present SteerAF, a novel automated alternative conformation prediction framework based on OpenFold ^36^ that leverages AlphaFold2’s distogram outputs. At inference time, our method uses the alternative signatures within the default conformation’s distogram to modulate the MSA input embeddings. Through rounds of MSA feature updates, SteerAF successfully sampled alternative conformations across multiple systems in the benchmarking datasets, including proteins with domain motions, fold-switching proteins, and transporters, while observing performance gains over most existing methods. Crucially, compared with computationally expensive MSA-subsampling or ensemble-sampling methods, our approach yields the desired conformations within an average of 10–20 runs. This efficiency strikes a favorable balance between prediction quality and runtime, satisfying the requirements of most practical alternative conformation prediction scenarios.

Furthermore, to address scenarios where reference experimental structures are unavailable, we implemented an effective reference-free state-selection strategy. Our strategy recovers the correct alternative conformation cluster in 85% (39/46) of the tested systems. We also demonstrate two downstream applications of SteerAF. First, the MSA-feature modifications pinpoint biologically meaningful residues, recovering experimentally characterized functional sites at ∼50% precision and a recall above 50% across two tested protein systems, far exceeding the 3.4% precision expected by chance. Second, seeding MD simulations of MdfA with predicted conformations reveals a multi-basin landscape that expands the sampled conformational space by more than threefold. This significantly complements structure prediction sampling and effectively aids in selecting the optimal prediction model. Overall, our framework provides an effective self-supervised approach for extracting alternative conformational states from the distogram information encoded in deep MSAs.

## 2 Results

### 2.1 Overview of SteerAF

Our inference-time guidance procedure (Figure 1A) is based on the paradigm of hallucination that was introduced in protein design tasks. ^37,38^ Hallucination is a reverse optimization method for de novo protein design based on PSP models. In this paradigm, the parameters of the PSP model are kept fixed, and the optimization target is not the model itself but the input amino acid sequence or MSA representation. Starting from a randomly initialized sequence or MSA input, Monte Carlo sampling, gradient-based optimization, or other sequence search strategies are used to iteratively update the input, so that the predicted structure becomes more confident and self-consistent. ^37–39^ Formally, let *M* denote the sequence or MSA input to be designed, and let *f*_0_ be a PSP model with fixed parameters θ. The hallucination process can be formulated as Equation (1):

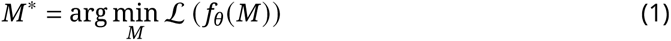

**Figure 1:**
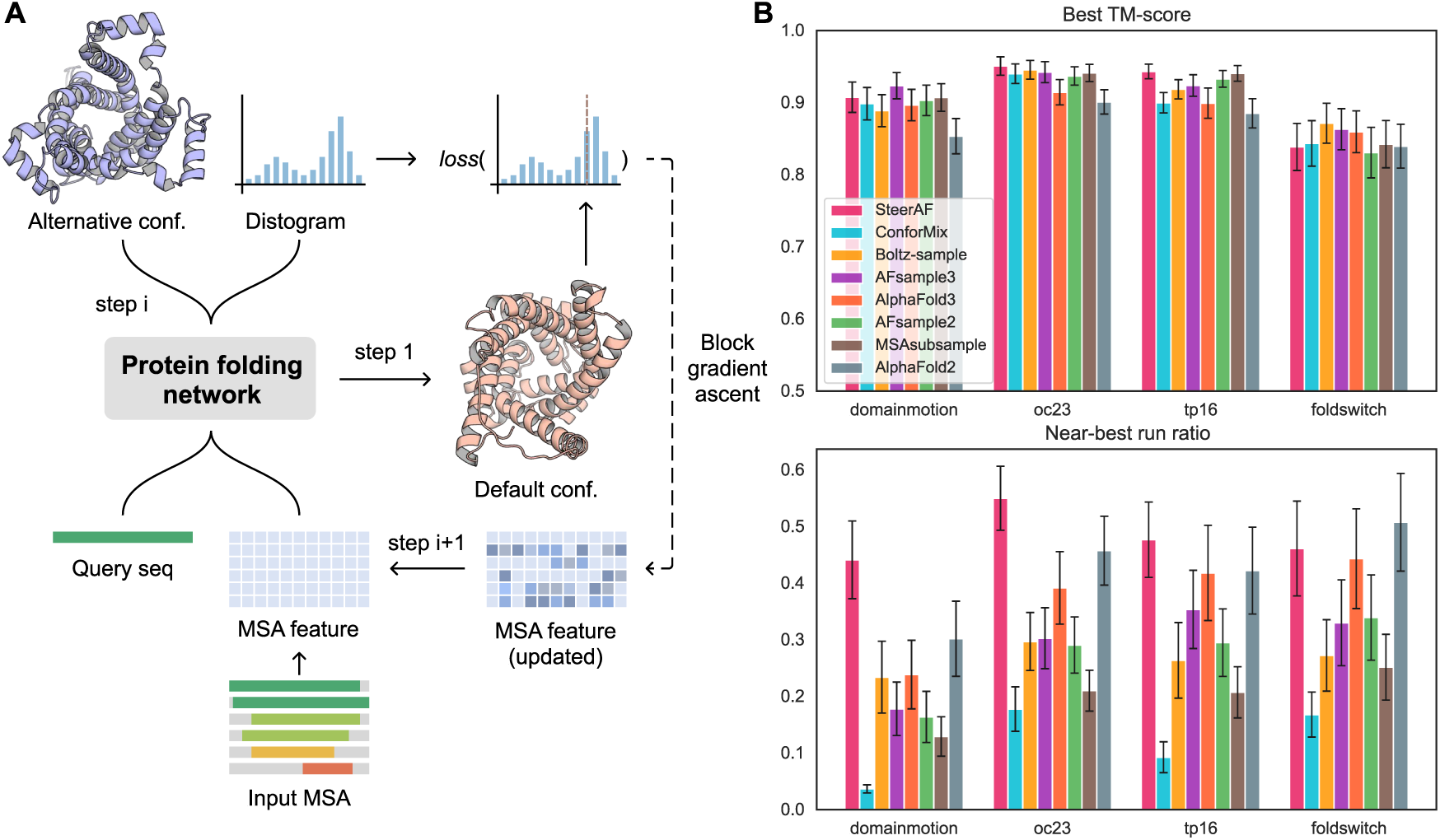
Overview of the SteerAF workflow. (A) The flow chart illustration of how SteerAF works, where the abbreviation “Conf” denotes conformation. (B) TM-score and near-best run ratio of SteerAF and other methods across four benchmark datasets; metric definitions are given in Methods Section 5.4. The SPF1 system was excluded from the benchmark due to its intractable length, which causes an out-of-memory issue.

where L is a task-specific loss function constructed from the predicted structural features. Since hallucination does not require a predefined target backbone and can flexibly incorporate additional design constraints, it has been widely used as a general framework for sequence-structure co-design ^39–41^ and experimental restraints-guided structure determination. ^31,42^

For PSP models applied to alternative conformation prediction, we make three assumptions (which have been mostly validated by our and others previous studies): (1) the high-probability distance bins of the distogram (i.e., the dominant peak) correspond to, or are highly correlated with, the actual pairwise distances in the protein structure model that the model ultimately outputs; (2) the MSA that guides the model toward alternative conformations differs from the MSA that directs it toward the default conformation; and (3) the PSP model itself is sufficiently powerful and already possesses knowledge of both the default and alternative conformations from the MSA, yielding multi-peak signals in the distogram that represent a mixture distribution of multiple states.

Apart from the *a prior* theory on co-evolutionary analysis explaining that MSA can carry footprints of multiple conformational states, ^32,33^ all these assumptions rely on the training objective of the PSP model’s distogram head. As shown in Equation (2), for each residue pair *k*, the training objective of the distogram minimizes the expected cross-entropy between the predicted distogram derived from the model output *z* given MSA inputs and the one-hot encoding of the true distance label from target structures *x* following the data distribution (*x* ∼ *p*_data_).

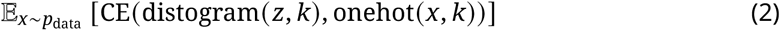

During training, the PSP model may have seen multiple possible structures for a target protein system, and the diversity increases further when homologous systems are included. ^1^ This turns the per-residue-pair training objective into a distribution-matching problem: the model must fit the empirical distribution of distance labels rather than a one-hot target. Since the model jointly optimizes the three-dimensional coordinates through a regression loss, ^1^ the distogram is encouraged to be strongly correlated with the coordinate output in each model rollout, helping to minimize the total loss. ^43^

Thus, we reasoned that using the alternative conformation signatures in the distogram to perform hallucination can yield an MSA that biases the model toward alternative conformations and allows the desired alternative conformation to be sampled. Specifically: (1) we freeze the model weights and set the numerical MSA input (the MSA feature) as the optimization target; (2) we feed the initial MSA feature into the model to obtain the default conformation; (3) we compute the cross-entropy (the distogram loss) between the pairwise distances of the default conformation and the pairwise distogram output by the model at each step, which serves as the core of the loss function, augmented with pLDDT as additional constraints. Our goal is to prevent the dominant peak of the model’s output distogram from overlapping too closely with the pairwise distances of the default conformation, while amplifying the signal of the alternative peak in the distogram as much as possible. (4) We backpropagate the loss gradient to the MSA feature, feed the updated MSA feature back into the model, and obtain the updated predicted conformation. (5) We repeat steps 3 and 4 multiple times to obtain a series of predicted conformations. Details of SteerAF algorithm are given in the Methods Sections 5.1 to 5.3. During the gradient update, we use several techniques (Method Section 5.3) that exploit the sparsity of the MSA-feature gradient.

### 2.2 Benchmark

We evaluated SteerAF on four datasets commonly used in the multi-conformation prediction field (Table 1), and compared it with other methods, including the AFsample series, ^11,44^ ConforMix, ^21^ Boltz-sample, ^45^ and the AlphaFold2/3 baselines. Prediction number and hyperparameter settings for SteerAF and the comparison methods are set as shown in Table 2. Detailed definitions and the rationale for choosing suitable values for SteerAF hyperparameters are given in Methods Sections 5.1 to 5.3 and 5.5. Because ConforMix requires predictions under multiple hyperparameter settings, with perturbations at different RMSD values, we followed the prediction count specified in its original paper. ^21^ To ensure a relatively fair comparison with baseline methods that are not specifically optimized for alternative conformation prediction, we set the number of AlphaFold2/3 baseline predictions to 1000.

**Table 1:** Benchmark datasets.

| Dataset | No. of systems | TM-score (mean / max / min) |
| --- | --- | --- |
| Domain motion<sup>20</sup> | 16 | 0.6813 / 0.8370 / 0.5070 |
| Open-close conformation (oc23)<sup>11</sup> | 23 | 0.5009 / 0.6450 / 0.3790 |
| Transmembrane (tp16)<sup>11</sup> | 15 | 0.7892 / 0.9690 / 0.3730 |
| Fold switch<sup>9,21</sup> | 15 | 0.6672 / 0.9004 / 0.4817 |

**Table 2:** Prediction-count, hyperparameter, and runtime settings per method.

| Method | Setting | Total models | Wall-clock hour mean (min-max) |
| --- | --- | --- | --- |
| SteerAF | $\rho \in \{10\%, 5\%\}; \text{lr} \in \{2 \times 10^{-2}, 2 \times 10^{-3}\}$.<br>10 runs for $\rho = 10\%$; 20 runs for $\rho = 5\%$. | **420**<br>(60 runs<br>$\times 7$ steps) | 1.35 (0.17–5.50) |
| ConforMix | 40 RMSD settings, spanning 0.5–20 Å;<br>20 models per run. | 800 | 2.73 (0.52–8.88) |
| Boltz-sample | $\beta \in \{\pm 0.15, \pm 0.30, \pm 0.45, \pm 0.60, \pm 0.75\}$;<br>50 models per $\beta$. | 500 | 0.42 (0.09–1.28) |
| AFsample3 | `msa_rand_fraction` = 0.4. | 500 | 0.47 (0.11–1.30) |
| AlphaFold3 | Repository defaults. | 1000 | 1.20 (0.43–3.42) |
| AFsample2 | `msa_rand_fraction` = 0.2. | 500 | 3.65 (0.42–13.98) |
| MSAsubsample<sup>7</sup> | max extra MSAs $\in$<br>{16, 32, 64, 128, 256, 512, 1024, 5120}. | 500 | 2.15 (0.22–8.27) |
| AlphaFold2 | Repository defaults. | 1000 | 8.58 (1.10–28.70) |

We compared the methods using five metrics: the TM-score and RMSD of the prediction closest to the experimental conformation (best TM-score/RMSD), the fraction of predicted structures close to the experimental conformation (fill ratio), the dispersion among such structures (cluster spread), and the near-best run ratio. Exact metric definitions are given in Methods Section 5.4. Best TM-score/RMSD measures each method’s absolute predictive ability, i.e., whether the alternative conformation can be predicted at all. The latter three metrics assess whether, in the absence of an experimental reference, a method can recover candidate alternative-conformation clusters from its predictions in an unsupervised manner. Specifically, a higher fill ratio, a smaller cluster spread, and a higher near-best run ratio indicate a denser alternative-conformation cluster that is easier to identify without supervision. The next section also discusses the effect of the near-best run ratio on sampling efficiency.

Figure 1B summarizes the TM-score and near-best run ratio of SteerAF across the four datasets, and more detailed results are reported in Figures S1 and S2. The results show that, at comparable prediction counts, SteerAF performs competitively with the best competing methods across most systems, for both MSA-prompting-based methods and methods that bias the diffusion process of generative models. On the fold-switch dataset, however, SteerAF is overall inferior to the diffusion-based prediction methods, Boltz-sample and AFsample3.

### 2.3 Sampling efficiency

When evaluating the capability of multi-conformation prediction algorithms, sampling efficiency is also worth considering. For example, the AFsample3 paper reports that their method can sample reasonably good alternative conformations within about 300 samples for most systems, although a few systems still require more, e.g., 780 samples for accurate prediction. ^44^ In our method, the number of protein structures predicted by the AF2 model depends simultaneously on two factors, the number of optimization steps per sample and the number of samples. The metric used to assess sampling efficiency should therefore be adpapted accordingly.

Specifically, within a single sampling run, we obtain a series of optimization results: in the first few steps, the result is usually closer to the default conformation, and only in the last or middle steps does the conformation approach the alternative conformation. This resembles the denoising process of a diffusion model, in that we do not obtain the desired result from the very beginning. In addition, other multi-conformation prediction studies have also noted that some predictions may correspond to transition-pathway conformations or intermediate conformations, ^8,11,46^ an observation supported by our predictions for systems such as P71447, P31133, and P33284 (UniProts). For both reasons, counting the absolute number of predictions similar to the alternative conformation may be inappropriate for a method like ours, which outputs a series of trajectories. We therefore count the number of trajectories that contain a near-best prediction (the number of trajectories equals the number of samples/runs) and compute the fraction of such trajectories among all runs, which we denote as the near-best run ratio. The near-best run ratio of other methods can be computed analogously, except that for those methods, we use the number of runs (i.e., the number of predicted structures) directly.

The results show that, when a prediction reaching 97% of the best TM-score is defined as near-best, on average 55.1% of the runs of our method sample an alternative conformation, reaching close to 100% for some systems. As shown in Figure 2A, compared with the three latest competing methods, SteerAF achieves a higher near-best run ratio while maintaining a comparable best TM-score. The low failure rate reduces the risk that candidate selection is misled by spurious predictions, enabling reliable identification of alternative conformations without experimental structures. This robustness, in turn, allows us to infer plausible point-mutation sites directly from the MSA optimization trajectory, offering new biological insights. SteerAF also requires only a relatively small number of runs and a few optimization steps per run. Under the same hyperparameter combination, most systems reached relatively saturated prediction performance within 10–20 runs, corresponding to about 70–140 predicted structures. Increasing the number of optimization steps improves the accuracy of alternative conformation prediction, but also lowers the fraction of desired structures among all predictions, making selection harder when no experimental alternative conformation is available. As shown in Figure 2C, the increase in the best TM-score slows markedly after 6 optimization steps, whereas the near-best prediction ratio continues to decline. Varying the number of runs shows a similar tradeoff. We therefore chose 6 optimization steps and 10 or 20 runs per hyperparameter combination as the balanced fixed settings for prediction.

**Figure 2:**
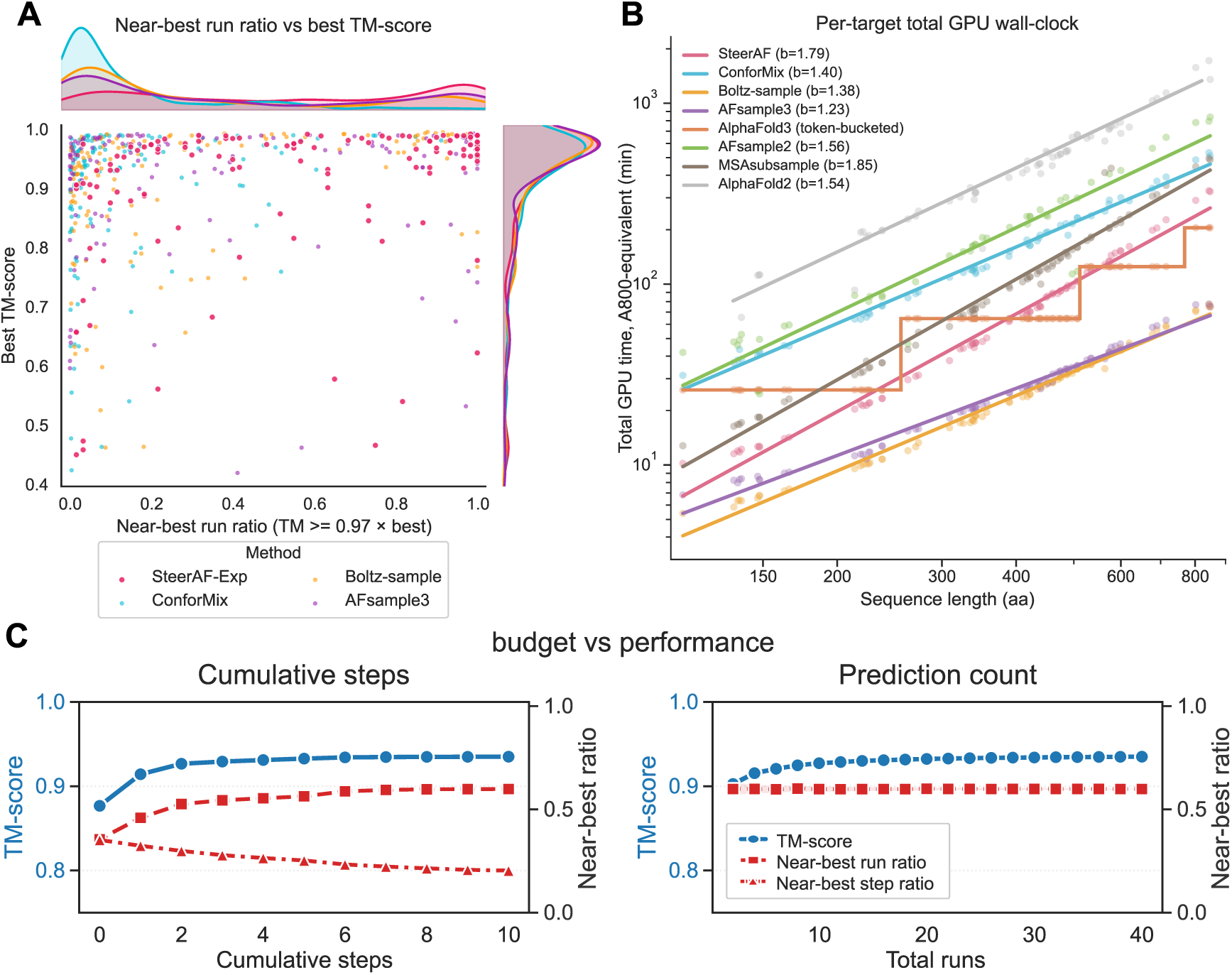
Sampling efficiency of SteerAF. (A) Near-best run ratio vs best TM-score distribution across all systems with four latest benchmark methods. (B) A800-equivalent total GPU time required to predict all structures for one target. AlphaFold3 uses a padding bucket of 256 / 512 / 768 / 1024 for better GPU utilization, so its runtime is displayed as a step function. (C) Best TM-score and near-best run/prediction ratio changes with number of optimization steps or predicting runs. In the first panel, the metric values are computed cumulatively over the optimization step and all preceding steps.

We also measured the GPU runtime of each method. Following the normalization procedure described in Method Section 5.6, runtimes measured on different hardware were converted to the A800-equivalent total GPU time required to predict all structures for one target, and the normalized runtime is shown in Table 2 and Figure 2B. AFsample3 and Boltz-sample, both based on diffusion-model sampling, were substantially faster than the other methods. SteerAF and default AlphaFold3 formed the next tier, whereas AlphaFold2-based baselines, which require more computation with MSA representation and Evoformer, were slower. SteerAF therefore retains a practical runtime advantage by using fewer total predicted structures than most baselines.

### 2.4 Reference-free conformation selection

For an alternative conformation prediction method, an important measure of practical utility is whether it can select a suitable alternative conformation candidate from all predictions in an unsupervised manner, in the absence of a resolved alternative conformation. AFsample3 and CF-random select reference-free candidates primarily by computing distance or Foldseek similarity matrices across all predicted structures, followed by principal component analysis (PCA) and clustering. ^9,44^ Inspired by this strategy, we devised a similarly intuitive and simple reference-free selection method suited to SteerAF.

We first identify, from the step-0 results (i.e., the default conformation), the common residue intervals with ordered secondary structure. We then compute, for every prediction restricted to these residues, the internal distance matrix between residues. If the common secondary-structure (SS) indices have length *l*, we obtain *N* internal distance matrices, forming an array of shape *N* × *l* × *l*. After flattening the last two dimensions of this array, we perform PCA to observe the main trends of structural variation across the predicted structures, and apply HDBSCAN ^47^ to obtain unsupervised conformational clusters. An illustration of this pipeline is shown in Figure 3A.

**Figure 3:**
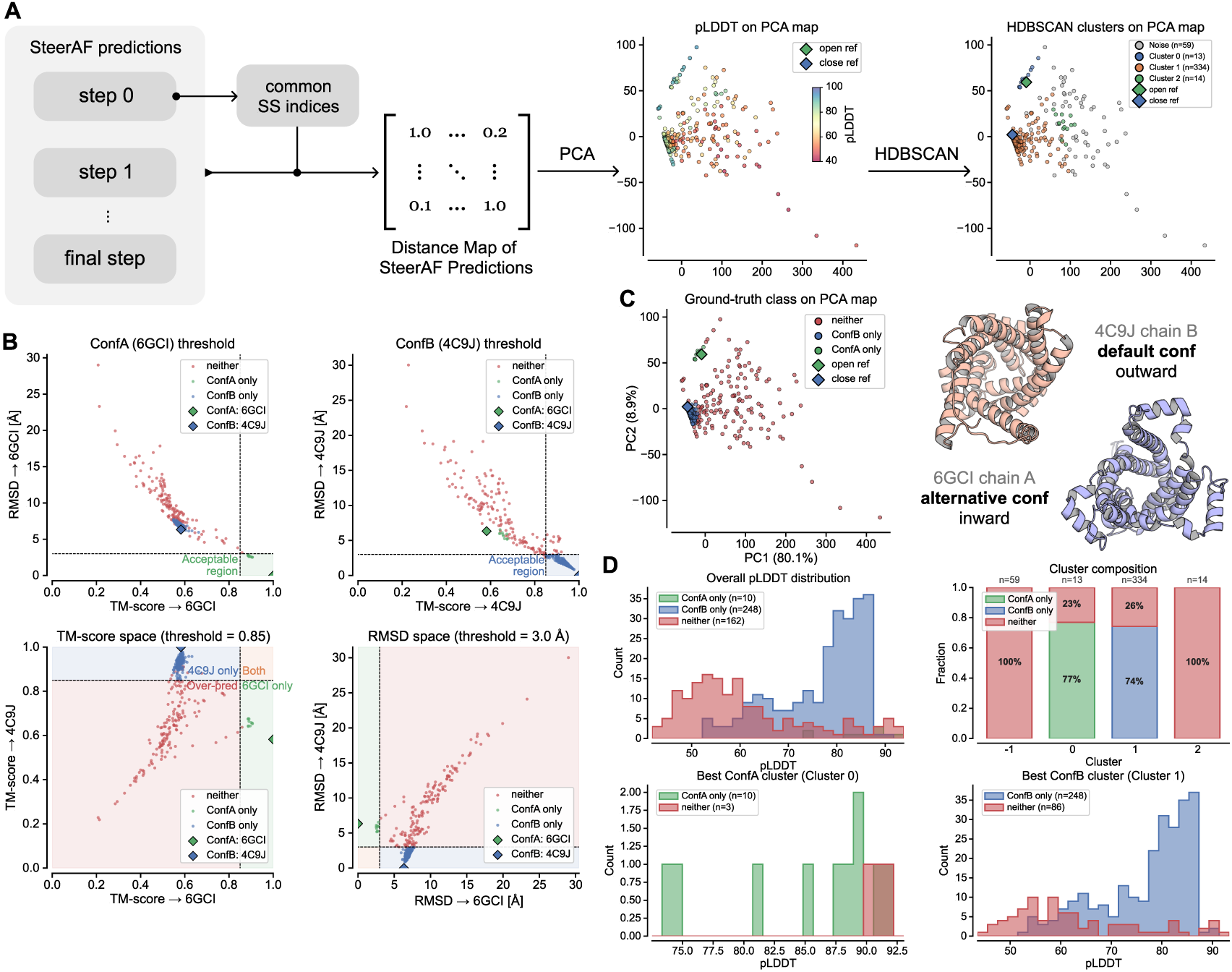
Reference-free conformational selection, illustrated using AAC3_OUT as an example. (A) PCA + HDBSCAN reference-free selection pipeline. The first PCA map is colored by pLDDT to guide the choice of appropriate HDBSCAN clustering parameters and the clustering itself, and the second is colored by the resulting HDBSCAN cluster assignment for AAC3_OUT. (B) Threshold determination from reference conformations for AAC3_OUT, where ConfA corresponds to PDB 6GCI and ConfB to PDB 4C9J. (C) Structures of the two reference conformations, with predicted structures on the PCA map colored according to which reference conformation (ConfA or ConfB) each is closer to, based on the threshold defined in (B). (D) Per-cluster composition of different conformations, and pLDDT distributions across all predictions (regardless of cluster) and within the representative clusters of ConfA and ConfB.

Below we show the performance of unsupervised conformational selection on the predictions for a mitochondrial ADP/ATP carrier (a member of the SLC25 mitochondrial carrier family), AAC3_OUT. To evaluate the selection, we first used the reference conformations to define a scoring criterion: a structure with TM-score > 0.85 and RMSD < 3 Å to a reference conformation is regarded as correctly predicting that conformation; the choice of these thresholds is justified in Figure 3B, and the predicted structures assigned to a reference conformation based on this threshold are shown in Figure 3C. This threshold may vary across protein systems: for example, GPCR conformations can be separated by relatively small structural differences and may require stricter thresholds, whereas many domain-motion proteins involve larger conformational changes and can tolerate looser criteria. We then applied the PCA + HDBSCAN pipeline described above without using any reference conformation. As shown in Figure 3D, cluster 0 captured more than 99% of the alternative conformations. The structures in cluster 0 that were not classified as acceptable alternative-conformation predictions were nevertheless highly similar to the alternative conformation, and cluster 0 contained no default conformations. Cluster 1, in contrast, captured the majority of the default conformations, with both default and alternative conformations accounting for more than 74% of their corresponding clusters. Thus, our clustering method achieves effective unsupervised alternative-conformation selection on AAC3_OUT. We also tested the selection method on additional protein systems. Here, a system is counted as a success only when both conformations are recovered as distinct clusters in the unsupervised pipeline (i.e., among the systems in which both conformations were successfully predicted, each conformation is clustered out, meaning it accounts for more than 50% of a corresponding cluster); the approach succeeded for 85% of systems (39/46), with most failures occurring in systems where the near-best run ratio was extremely low. For reference, CF-random reports an 81% success rate (26/32) in its reference-free (blind mode) selection on its in-house fold-switch dataset, ^9^ while AFsample3 scores success per conformation rather than per protein, achieving a top-1 success rate of 77% and a top-10 success rate of 89% across 238 conformations. ^44^

### 2.5 Applications of SteerAF

#### 2.5.1 Interpretability and biological evidence in the modified MSA feature

A key design choice in SteerAF is to optimize the MSA input feature rather than intermediate network representations. This allows us to interpret the resulting residue-level modifications directly as biologically meaningful guidance for experiment design. Here we take PGK1 (human phosphoglycerate kinase, UniProt P00558) as an example to illustrate the relationship between SteerAF’s modifications to the MSA feature and known structurally or functionally important sites (Figure 4A).

**Figure 4:**
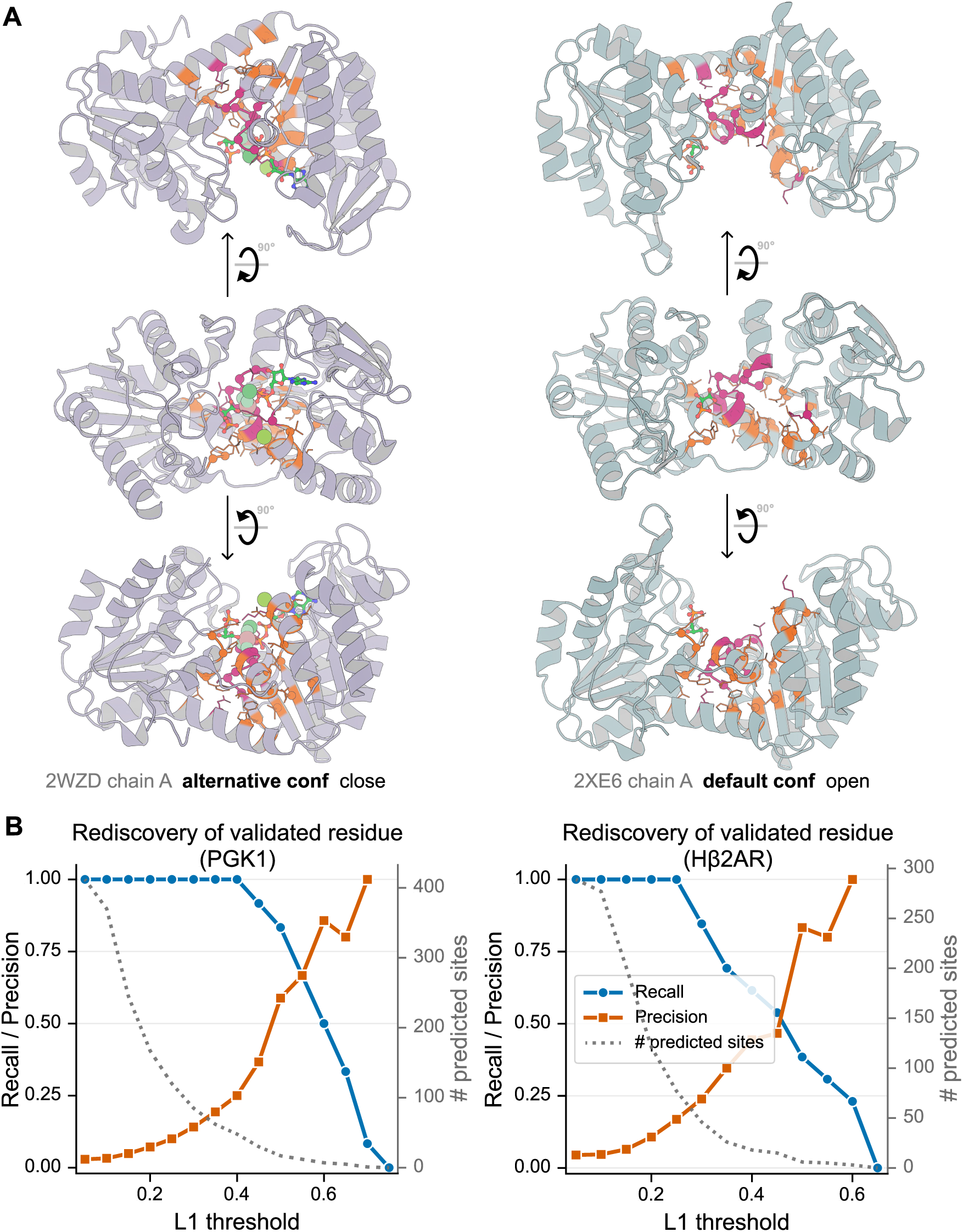
Biological interpretability of SteerAF MSA-feature modifications. (A) Relationship between the important sites identified by PSSM analysis and the structurally or functionally important residues reported in the literature (Method Section 5.7), using PGK1 (UniProt P00558) as an example. These literature-reported residues are marked in red, and residue sites inferred from PSSM analysis, defined as positions where the L1-norm of the per-residue amino-acid distribution in the PSSM exceeds 0.4, are marked in orange. (B) Important sites of PGK1 and hβ_2_AR are selected by varying the threshold on the L1-norm of the per-residue amino-acid distribution in the PSSM modification, and the resulting precision and recall are computed.

We collected studies published since 2000 that reported residues important for PGK1 conformational transitions or function-conformation coupling. Detailed sources are provided in Method Section 5.7. The curated sites fall into four groups, the first three of which have direct biological experimental support:

1. closure-trigger and interdomain-communication residues: R38, K219, N336, and E343;
2. main-hinge and beta-strand-L regulatory residues: F165, E192, F196, S392, and T393;
3. open-state binding and fully closed/TSA-like state regulatory residues: D218 and D374;
4. computationally predicted allosteric nodes: G371, G372, and G394.

We then analyzed the MSA features from near-best runs, defined here as runs containing predictions that reached at least 97% of the best TM-score for the target. For every site and every alignment sequence, we independently normalized the feature vector and treated it as an amino-acid probability distribution. Averaging these distributions across all alignments and all selected MSA features yielded a position-specific scoring matrix (PSSM) for the full sequence. We computed the PSSM for the last near-best step in each near-best run, the PSSM for the initial step-0 prediction, and their difference. The full visualizations are provided in the Figure S3. The difference between the alternative conformation PSSM and the step-0 PSSM shows that sites with substantial residue-level changes are highly site-specific. We then applied a range of thresholds to select modified sites, compared the selected candidate sites with the curated functional residues, and computed precision and recall. As shown in Figure 4B, across a broad threshold range of 0.3–0.5, SteerAF maintains a precision of 15%–50% and a recall above 60%, far exceeding the 3.4% (14/416) precision expected from random selection.

We further performed a site-restoration test as an orthogonal validation of the PSSM analysis. For sites experimentally or computationally implicated in PGK1 conformational regulation, we restored the last near-best step MSA feature values to their corresponding step-0 values. As a control, we randomly selected twice as many sites outside this curated set that nevertheless had high or comparable MSA-feature modification levels, and restored those sites in the same way. We then fed both restored MSA features back into AlphaFold2. If restoring the curated sites, despite involving only half as many sites as the control set, caused a larger decrease in TM-score to the alternative conformation, these sites would be more important for SteerAF-mediated alternative conformation prediction. Figure S4 shows that the median and 75th percentile of TM-score after curated-site restoration were both lower than those after control-site restoration, which is consistent with the expectation.

As a supplement, we performed a similar analysis on the hβ_2_AR receptor and plotted the relationship between selection threshold, precision, and recall in the same format as PGK1, and the result is also shown in Figure 4B. This additional analysis likewise shows that selecting mutational sites from SteerAF MSA-feature modifications is far more reliable than random selection. Together, the PSSM statistics and site-restoration test indicate that SteerAF predictions, and specifically the sites selected by SteerAF MSA-feature modification, are likely biologically relevant rather than arbitrary perturbations that merely maximize the distance from the default conformation.

#### 2.5.2 Leveraging predicted structures to seed conformational exploration in MD simulations

As a proof of concept, we asked whether SteerAF predictions can serve not only as static alternative conformation candidates, but also as starting points for molecular dynamics (MD) simulations that explore the nearby conformational landscape. ^48^ We tested this idea on MdfA, a bacterial multidrug/H^+^ antiporter, for which our reference-free selection strategy failed to identify the alternative conformation candidates because all predictions near the alternative conformation were classified as outliers. Rather than initiating MD from a single predicted structure, we treated the SteerAF prediction set as a coarse map of conformational space. From the 420 predicted structures, we performed distance-based PCA and selected 50 representative structures such that the selected starting points covered both high-density regions and peripheral predicted conformations (Figure 5A).

**Figure 5:**
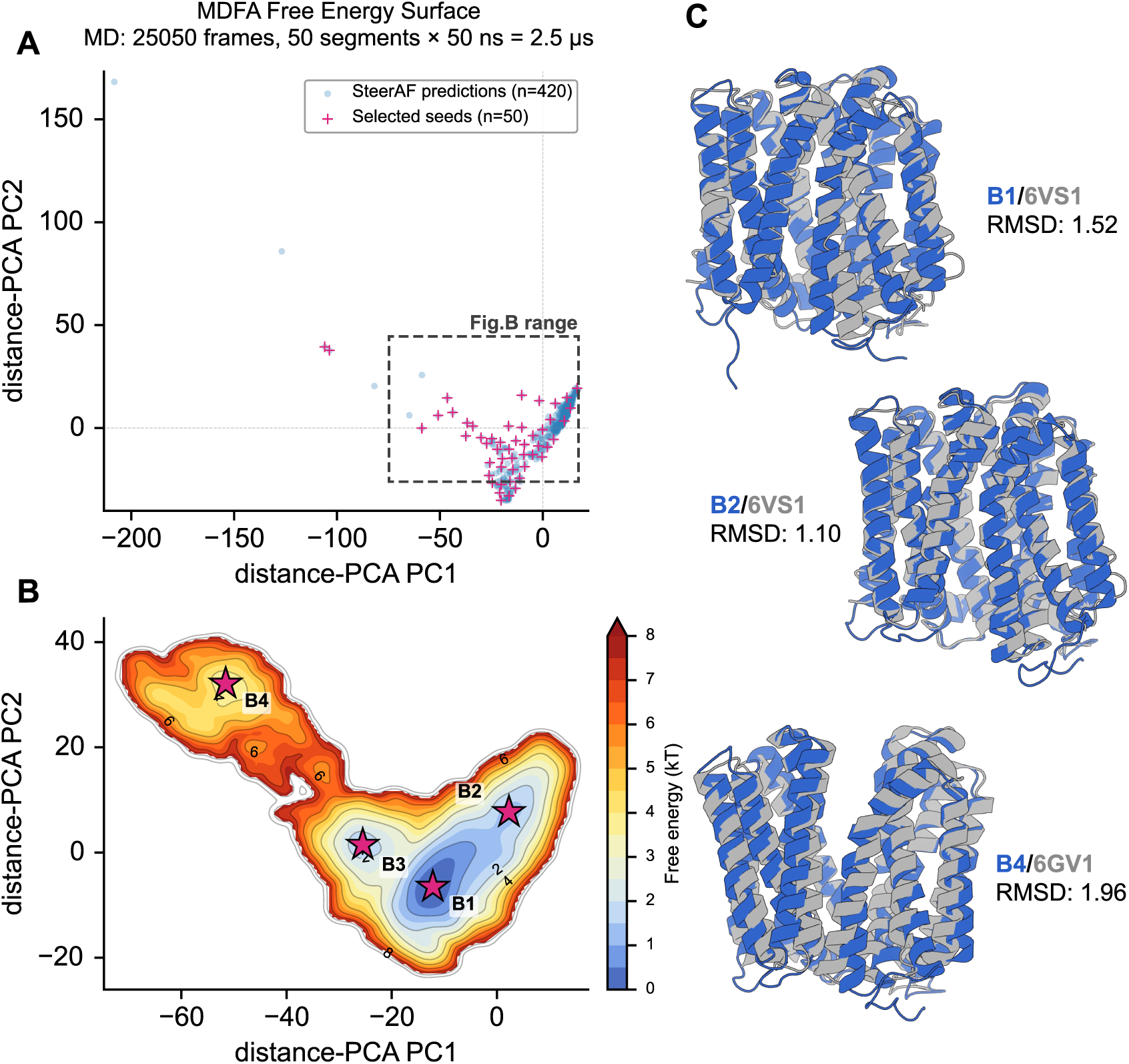
SteerAF predictions as starting points for MD-based free-energy exploration on MdfA. (A) Distance-PCA space of 420 SteerAF predictions (blue dots) and 50 selected representative structures (red cross). The 50 representatives were selected within this PC1–PC2 plane to cover the landscape evenly rather than over-sampling the densest region (Methods Section 5.8). (B) We use representative structures as relay targets in a cascaded targeted-MD protocol, yielding 2.5 μs of aggregate sampling. The resulting two-dimensional free-energy surface, estimated by kernel density estimation in the same PCA space, reveals four basins (B1–B4, red stars). (C) Basin structures and their distances to the reference experimental conformations. B3 may correspond to an occluded state without an experimentally resolved structure.

Starting from the densest selected prediction, we built a single MD system with the protein embedded in an explicit POPC bilayer. Using a relay targeted-MD protocol, we drove the system through the 50 selected structures in the prediction-derived PCA space. All 50 targeted-MD simulations converged to their assigned targets, with final RMSDs of 0.6–1.2 Å. The resulting structures were then used as starting points for 50 independent 50 ns production simulations, yielding a total of 2.5 µs of aggregate MD sampling (Methods Section 5.8). To compare the MD trajectories with the original prediction ensemble, we projected all production frames (excluding the first 10% of each trajectory) back onto the same distance-based PCA space and estimated a two-dimensional free-energy surface (Figure 5B). We note that this free-energy estimate is rough and that the sampling has not fully converged. Nonetheless, the MD frames covered a substantially broader local region than the static prediction ensemble: on a 2.0-PC grid spanning the MD-accessible region, the original predictions occupied 123 cells, whereas the MD trajectories occupied 437 cells, including 356 grid cells not visited by the prediction set.

The resulting conformational landscape contained four basins (Figure 5B). The global minimum was B1 at PC1 = -12.2 and PC2 = -6.5, followed by B2 at PC1 = 2.3 and PC2 = 7.7, B3 at PC1 = -25.5 and PC2 = 1.5, and B4 at PC1 = -51.6 and PC2 = 32.1. For each basin, we extracted the nearest trajectory frame, and the distances between the extracted frames and basin centers were small in PCA space (all < 0.1 PC units for B1–B4, respectively). Notably, the nearest original predicted structures and reference structure 6VS1 were close to B1 and B2 but much farther from B3 and B4, while reference structure 6GV1 was closer to B4 (Figure 5C). B3 may show as an intermediate occluded state, in line with the intermediate or occluded conformations of MdfA inferred from previous MD simulations and double electron–electron resonance (DEER) experiments. ^49–53^ Notably, predictions near B4 were not resolved as a distinct cluster by our reference-free selection strategy, yet MD sampling successfully recovered a well-defined free-energy basin at this location. The same workflow was also applied to the mitochondrial ADP/ATP carrier AAC3 (UniProt P18238), where it likewise expanded the sampled conformational space and resolved a multi-basin free-energy landscape (Figure S5).

Together, these observations indicate that MD relaxation and short production sampling can refine and extend the conformational information provided by SteerAF predictions. These results suggest a practical workflow in which SteerAF rapidly proposes diverse, model-informed conformational seeds, and MD simulations then convert these seeds into a physically relaxed ensemble suitable for conformational landscape analysis. This approach also identifies free energy basins, thereby facilitating the selection of alternative and metastable conformations.

## 3 Discussion

Despite achieving competitive performance on two-state conformational benchmarks, our method performs relatively poorly on the fold-switch dataset. We suspect that, because our method relies on gradient descent in a continuous optimization space and the distograms of two distinct fold-switch conformations often differ drastically, the alternative-peak signal encoded in the default-conformation distogram may not provide sufficient guidance for reaching the alternative state. As a result, distogram-based gradient descent struggles to generate accurate predictions for fold-switch proteins. Furthermore, other AF2-based methods also perform poorly on this dataset (Figure 1B and Figure S1), suggesting that these limitations may extend beyond our specific optimization strategy and may partly reflect inherent constraints of the underlying AF2 model or the quality of the fold-switching dataset. ^9^

SteerAF also performs poorly on proteins that adopt three or more distinct conformations, such as multiple terminal states of domain motions, allosteric endpoints, and transporter channel states. These multi-discrete states are distinct from intermediate or transition-state conformations along continuous conformational trajectories. We tested it on an in-house multi-conformation protein dataset; the results show that 7/18 proteins yielded only one predicted conformation, 7/18 proteins yielded two, and 2/18 proteins yielded three or more. We reasoned that the alternative peaks from multiple conformations are mixed within a single distogram, whereas our method relies on driving the predicted conformation’s distogram away from the default conformation’s distribution; this loss function inherently cannot separate the alternative peak signals originating from different conformations. AlphaFold2 was likewise not specifically trained to disentangle different possible signals in the distance representation, so the model’s capacity during gradient backpropagation may also be insufficient to disentangle the mixed alternative-conformation signals, leading to a failure to find the correct MSA optimization direction. We anticipate a hybrid approach of differentiable MSA prompting and hallucination to solve these multi-discrete-state systems as well as fold-switching systems in the future.

Additionally, our method underperformed relative to others on only two systems: A6UVT1 and PF0708. For A6UVT1, neither AlphaFold2 nor AF2-based methods predicted the alternative conformation, whereas AlphaFold3, AF3-based methods, and Boltz-based methods all succeeded. All PDB depositions of A6UVT1 date from 2019, whereas the training data cutoff for AlphaFold2 is 2018-04-30^1^ and for AlphaFold3 and Boltz is in or after 2021. ^54–56^ Therefore, we cannot exclude the possibility of training-data leakage for the methods that successfully predicted this alternative conformation. For PF0708, SteerAF succeeded at a 20% sample ratio. We accordingly recommend that users applying SteerAF to entirely new systems explore a wider hyperparameter range, with a sample ratio ρ ≤ 20% and learning rate lr ≤ 2e-2 as a reasonable range.

In future work, we plan to extend our framework to all-atom structure prediction models. BoltzDesign1^41^ has already demonstrated that hallucination can be effectively performed on a diffusion-based AlphaFold3-like all-atom model ^54^ by backpropagating exclusively through its Pairformer, Confidence, and Distogram modules, while the diffusion process is completely bypassed. In the design context, sequence optimization is driven by both confidence-based and distogram-based losses. For alternative state prediction, our framework applies a similar strategy to optimize the MSA features and shifts the emphasis toward distilling the signals encoded in the distogram. This direct relationship between the two tasks outlines a clear route for applying our approach to the prediction and design of multi-state systems. Since our framework operates at the MSA level to capture biologically meaningful modifications, which is different from contemporaneous related preprints, ^57,58^ and provides a straightforward yet effective strategy for selecting candidate alternative conformations without requiring experimental structures, we anticipate that it will enable an interpretable and self-supervised route to predict and design multi-state biomolecular systems.

## 4 Conclusion

In this work, we demonstrated that the mixture of signatures from multiple conformations encoded in AF2’s predicted distogram can be effectively exploited by SteerAF, without updating the pretrained model weights or requiring large-scale sampling. By optimizing the MSA feature through gradient-based inference, SteerAF turns these signatures into effective guidance that steers AlphaFold2 from its dominant conformation toward alternative ones. Across diverse two-state systems, this strategy matches or surpasses existing methods while requiring far fewer samples, showing that exploiting a model’s intrinsic conformational knowledge offers an efficient yet high-quality route to alternative-conformation prediction. Crucially, because we optimize the interpretable MSA input rather than internal network representations, the resulting sparse, site-specific modifications can be cross-validated against experimentally characterized functional residues—indicating that SteerAF’s predictions are not arbitrary perturbations engineered to escape the default conformation, but instead carry genuine biological meaning, thereby extending a structure-prediction tool into a hypothesis-generating instrument that can inform experimental design. We further show that these predictions can support unsupervised conformational screening when no experimental reference is available, and can serve as starting points for MD simulations that explore the conformational landscape, positioning SteerAF not merely as a standalone predictor but as an efficient, interpretable entry point into real-world alternative-conformation discovery pipelines. Although limitations imposed by the underlying model persist for fold-switch proteins and systems with multiple discrete states, the principle of distogram guidance is not tied to any particular network architecture, and we anticipate extending it to more extensive all-atom structure-prediction models to reach an even broader range of biological systems for both structure prediction and design.

## 5 Method

### 5.1 Guided MSA optimization for alternative-conformation folding

The core algorithms of SteerAF are distogram-guided optimization and the block gradient ascent method used during optimization. The distogram-guided optimization pipeline is built on OpenFold (the open-source reproduction of the official AlphaFold2 project), with modifications focused on OpenFold’s inference mode. SteerAF is therefore essentially an inference-time optimization of OpenFold.

In the preparation stage before each round of SteerAF optimization, we provide the system information, i.e., the input sequence *s* of the protein system, the MSA corresponding to *s*, and the various hyperparameters required by SteerAF. In AlphaFold2’s design, the model takes three types of inputs: the input sequence, the MSA, and templates. Because our method does not use template information, only the first two are used in practice. For SteerAF’s hyperparameters, we define *T* as the number of distogram-guided optimization steps, and *K* as the number of initial steps during which AlphaFold2 recycling is not performed.

After AlphaFold’s default data-processing pipeline, the MSA is numerically encoded into the MSA feature and the extra MSA feature, whose sequence counts are typically in a 1:10 ratio. This split balances the use of as much evolutionary information as possible against computational cost and memory pressure. The MSA feature comprises the representative sequences of the MSA and is the input the model relies on most heavily during inference. Distogram-guided optimization therefore optimizes only the MSA feature, keeping the extra MSA feature fixed.

SteerAF then performs *T* optimization steps following the pipeline described in Algorithm 1 and in the Results section. The MSA feature input at step *i* is denoted as *M*_*i*_. From step *K* onward, the model performs one recycle during inference, where the MSA feature input for recycle 0 is *M*_0_ and that for recycle 1 is *M*_*i*_; the gradient update of *M*_*i*_ is performed via block gradient ascent. The last step performs no gradient backpropagation or updates, so SteerAF performs *T* + 1 predictions and outputs *T* + 1 predicted structures in total, of which the first *T* steps involve MSA-feature gradient updates.

#### Algorithm 1 Guided MSA Optimization

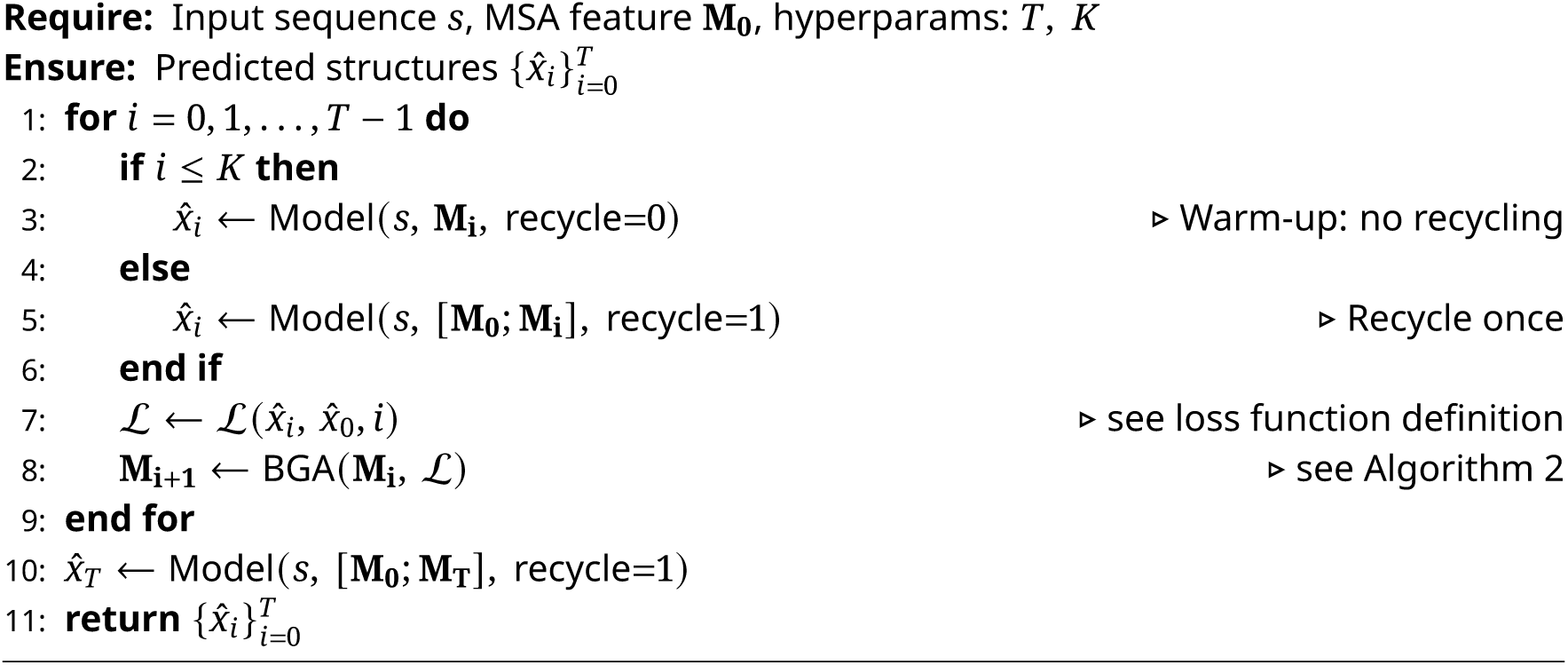

### 5.2 Loss function

Our loss function is a weighted sum of a distogram loss and a pLDDT loss, where γ_*i*_ is the weighting coefficient (Equation (3)). The pLDDT loss is introduced because, in the later stages of multi-step optimization, the gradient of the increasing cross-entropy loss does not decay to zero. Therefore, we add a pLDDT constraint to prevent the optimization from drifting away from the normal MSA distribution.

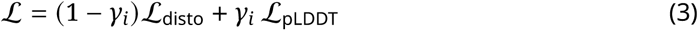

The distogram loss is computed in two stages, following the same cross-entropy form as AlphaFold’s official distogram loss but with an additional mask that upweights residue pairs whose step-0 distance bins are already poorly explained by the current distogram. We first compute the default distogram cross-entropy, treating the distance bins of the step-0 predicted structure (*x*_0_) as the target one-hot distribution and the distogram of the step-*i* prediction (*z*_*i*_) as the predicted distribution for each residue pair *k* (Equation (4)).

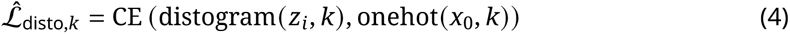

We then construct a square mask of the same shape as the *L* × *L* residue-pair matrix according to 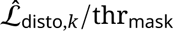. We apply it in a second distogram-loss calculation, with thr_mask_ = 4 by default. This masking strategy increases the weight of residue pairs whose step-0 distance bins and distogram have high cross-entropy, which are precisely the positions more likely to contain alternative-peak information (Equation (5)). At each optimization step, SteerAF performs gradient ascent in the direction of increasing this final loss.

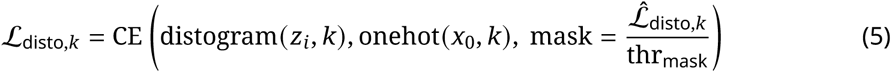

To keep the two loss terms on comparable scales, the pLDDT loss is rescaled (Equation (6)).

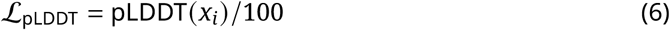

γ_*i*_is computed as follows. *K*_*p*_ denotes the step from which the pLDDT loss is added to the loss function, and λ_*p*_ is a fixed scaling coefficient for the pLDDT-loss weight, with a default of λ_*p*_ = 0.5. From step *K*_*p*_ onward, γ_*i*_ramps up linearly with the optimization step to its maximum value λ_*p*_ at step *T* (Equation (7)).

### 5.3 Block gradient ascent

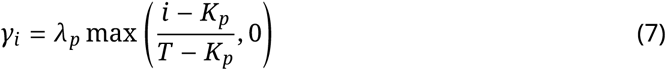

The MSA-feature modifications that drive AlphaFold2 to fold toward alternative conformations are sparse, which we confirmed experimentally. Compared with SteerAF-GA, which updates the entire MSA feature in the conventional way, SteerAF with sparse MSA-feature updates (the block gradient ascent, BGA, described in this section) improved the median best TM-score across all systems by 0.009, as shown in Figure S6. Relative to GA, BGA improved the TM-score by more than 0.05 on 7 systems and clearly worsened it on only 3, so we conclude that optimizing only a subset of MSA sites can match, or even slightly exceed, the performance of optimizing all MSA sites. Moreover, in the optimized MSA feature, the modifications made by SteerAF-BGA cover only a small subset of sites and are markedly site-specific. As noted for the interpretability question in Results Section 2.5.1, when predictive performance is comparable, we prefer the prediction scheme that provides more site-specific insight; therefore, except where SteerAF-GA is explicitly discussed in this section, all other parts of the paper discuss and present SteerAF-BGA results.

Block coordinate descent, or block gradient descent, is a variant of traditional coordinate descent. The main difference between traditional coordinate descent and gradient descent is that gradient descent updates the values of all variables after each backpropagation, whereas coordinate descent updates the value of only one variable (one coordinate) after each backpropagation. Correspondingly, block gradient descent updates the values of a group of variables (a block) after one backpropagation. The update order among blocks can be decided in several ways; here we choose an importance-sampling method based on the Gauss–Southwell selection rule, ^59,60^ in which the probability of selecting any given block depends on the magnitude of that block’s gradient, and a fixed proportion of blocks is drawn according to this probability for updating. We define a block as one residue (*s*, *r*) in the MSA feature *M*, where *n* is the row position of the residue in the MSA and *r* is the residue position relative to the input sequence; each block is a vector of length *D*, where *D* is the feature dimension of the MSA feature. The detailed BGA algorithm is given in Algorithm 2.

Note that, when updating the gradient, similar to the gradient regularization in ColabDesign, ^37,39,40^ we also regularize the gradient. We tried three regularization schemes in total: normalizing to a unit vector (unit), normalizing by the BGA sampling probability (exp), and normalizing by the original gradient norm (norm). The detailed comparison results are shown in Figures S6 and S7. These three schemes have slightly different strengths across systems, but show no significant difference in overall performance on the full dataset. Based on our current experimental comparison, SteerAF-exp alone is sufficient to recover the best performance among the three schemes for the vast majority of systems in the four datasets used here. Therefore, unless a specific regularization scheme is stated otherwise, all other results in this manuscript use SteerAF-exp.

#### Algorithm 2 BGA: Block Gradient Ascent with Importance Sampling

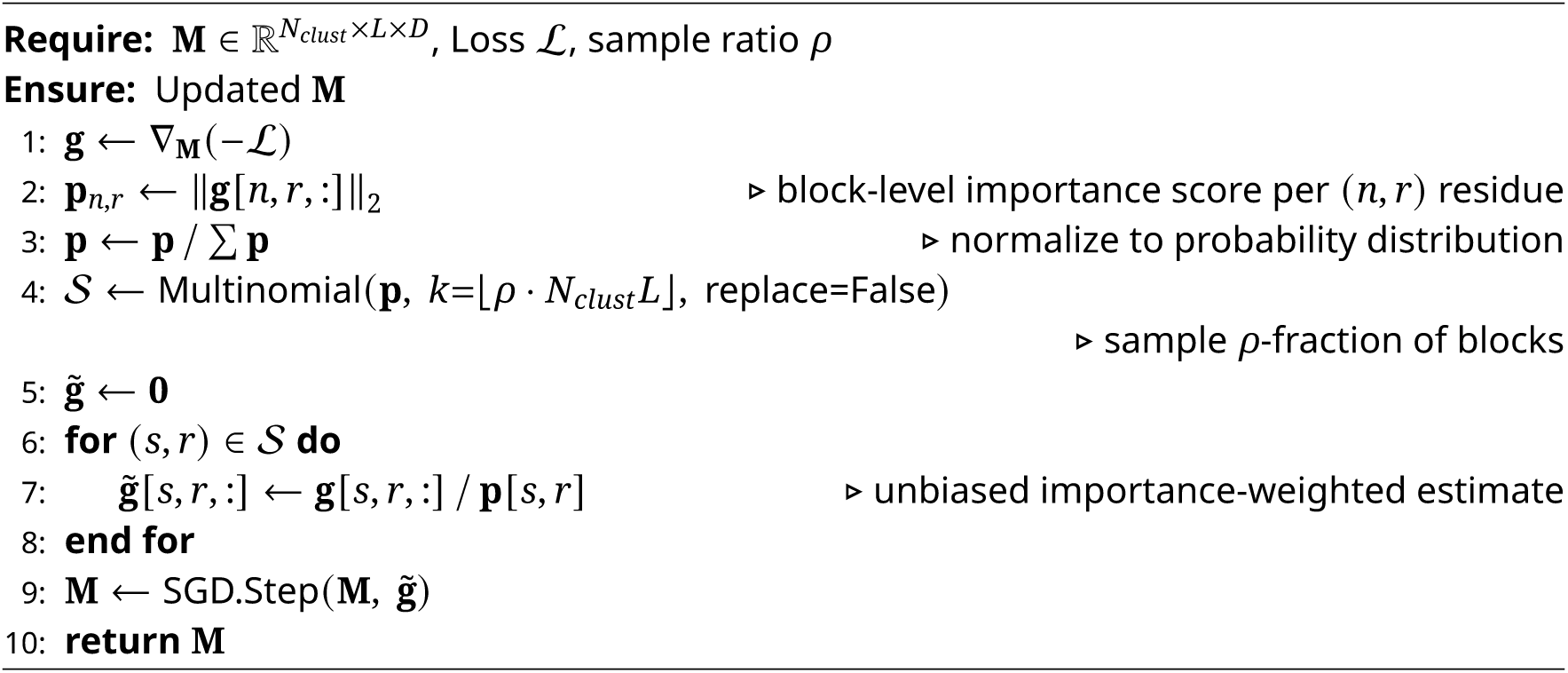

### 5.4 Metrics

#### Fill ratio

For each target conformation, we define acceptable TM-score and RMSD thresholds, denoted thr_tms_ and thr_RMSD_. A predicted structure is counted as successfully close to the target conformation when it satisfies both TM-score > thr_tms_ and RMSD < thr_RMSD_. If the number of such predictions is *N*_confX_ and the total number of predictions is *N*_total_, the fill ratio is defined as Equation (8):

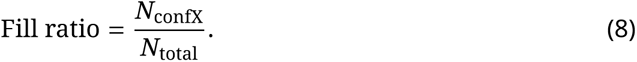

This metric captures sampling efficiency and is also important for reference-free state selection. A higher fill ratio provides clearer principal-component signals and makes target-like structures more likely to form a coherent cluster.

#### Cluster spread

For the *N*_confX_ structures close to the target conformation, we compute the product of the standard deviations of their TM-score and RMSD to the target (Equation (9)):

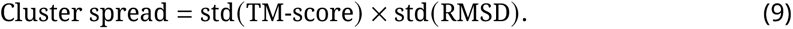

Cluster spread measures how concentrated the target-like predictions are. More concentrated predictions are easier to recover as a single cluster in reference-free analyses rather than being treated as scattered outliers.

#### Near-best run ratio

We use two related definitions of near-best run ratio in this manuscript. In the first definition, for a given target system, a prediction or run is considered near-best if it reaches at least 97% of the best TM-score achieved for that target. These are referred to as near-best predictions or near-best runs, respectively. The near-best run ratio is then defined as Equation (10):

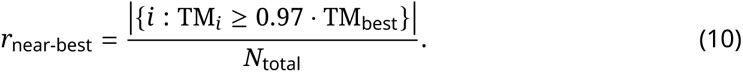

In the second definition, the same 97% criterion is retained, but near-best runs must also reach an absolute TM-score greater than 0.85 (Equation (11)):

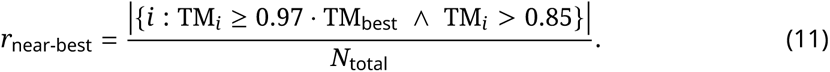

The first definition is used only in Figure 2, where the best TM-score is shown together with the near-best ratio. The second definition is used elsewhere because the near-best ratio is intended to measure how efficiently a method reaches the target conformation. Without an absolute TM-score requirement, a poorly predicted target could still have a high near-best ratio simply because many low-quality predictions are close to an equally low-quality best prediction. Thus, we use the second definition of near-best ratio when there is a lack of best TM-score/RMSD information.

### 5.5 Recommended hyperparameters

We tested three important SteerAF hyperparameters over a broad range: learning rate lr (2×10^−3^, 2×10^−2^, and 5×10^−2^), fixed pLDDT loss weight λ_*p*_ (0.33, 0.50, 0.67, and 0.80), and sample ratio ρ (5% and 10%). This hyperparameter tuning result was shown in Figure S8. At a sample ratio of 10%, all combinations of learning rate and pLDDT loss weight were evaluated on the OC23_TP16 and fold-switch datasets. The results showed no strong monotonic trend across the hyperparameter space, and overall performance was similar across many settings. However, the peaks of cross-system Gaussian-kernel-smoothed performance curves were more concentrated near a learning rate of 2×10^−2^ and a pLDDT loss weight of 0.50, suggesting that this combination is suitable for most systems.

Additional comparisons across sample ratios and learning rates showed that some systems are more sensitive to sample ratio, and that a better learning rate can improve difficult systems. However, larger sample ratios and learning rates also increase the risk of excessive gradient ascent, producing more low-quality predictions and more incorrectly or incompletely folded structures. To balance predictive performance with structural quality, we recommend λ_*p*_= 0.50, lr ∈ {2×10^−3^, 2×10^−2^ }, and ρ ∈ {5%, 10%} as practical default settings. More detailed recommended hyperparameter ranges are provided in Table S1.

### 5.6 Runtime normalization across different hardware

Because the methods were benchmarked on a heterogeneous pool of GPUs, all runtimes were converted to an A800-equivalent total GPU time per target. This quantity is defined as the time required to generate one sample or prediction on an A800, multiplied by the total number of predictions produced per target. GPU-to-GPU conversion factors were not assumed a priori; instead, they were estimated empirically by running the same target system with the same method on different GPUs. Conversion factors were applied only within the same method family, because relative slowdowns on older hardware depend on the compute kernels used by each method. Diffusion-based methods, for example, are more hardware-sensitive than AlphaFold2-based methods. Methods already run on A800 required no conversion. The normalized runtimes reported here are intended only for relative comparison across methods, not as absolute runtime guarantees. Raw runtimes, conversion factors, and their derivation are available in the source code.

### 5.7 Residue sites supported by experimental validation

For P00558 (human PGK1), we curated experimentally validated sites involved in open–closed conformational coupling, catalytic-state formation, or hinge regulation (Table S2). These references were independent of SteerAF predictions and used only to evaluate whether SteerAF MSA-feature modifications recover biologically meaningful residues.

The curated sites listed here do not exhaustively cover all functional positions; unannotated residues should therefore not be treated as irrelevant. Rather, this set represents a well-supported subset of residues with established roles in conformational regulation, which we used in the PSSM analysis to gauge the biological relevance of SteerAF MSA-feature modifications without designating unvalidated positions as negative examples. The same principle applies to the hβ_2_AR analysis below.

In addition to the experimentally validated residues above, G371, G372, and G394 were retained as computationally predicted allosteric nodes from force-distribution analysis. ^61^ These residues provide computational support for the allosteric pathway but were not treated as direct mutational evidence in the same sense as the experimentally validated sites.

For P07550 (human β_2_AR), we curated a set of residues involved in active-state conformational selection, drawing on direct mutational evidence for constitutive activation (Table S3). Beyond the directly mutationally validated sites above, we additionally retained I121, R131, Y132, P211, W286, N318, N322, P323, and Y326 in the PSSM analysis as structurally characterized microswitch positions. These residues correspond to established hβ_2_AR microswitch motifs—the DRY motif (D130/R131/Y132), the PIF/connector motif (P211/I121/F282), the rotamer toggle-switch region (C285/W286), and the NPxxY motif (N322/P323/Y326)—identified through active- versus inactive-state structural comparisons and allosteric network analyses. ^62^ N318 (N7.45) was additionally included as a structural packing node: comparison of the inactive- and nanobody-stabilized active-state structures reveals that the I121/F282/N318 packing arrangement rearranges upon receptor activation, coupling TM6 outward rotation to repositioning of TM7. ^63^

### 5.8 MD simulations for conformational exploration

To turn a SteerAF prediction set into a physically relaxed conformational ensemble, we used a general all-atom MD protocol that treats the predicted structures as a coarse map of conformational space. The protocol below is system-agnostic, and the same settings were applied across systems.

#### Seed selection

The predicted ensemble was first embedded in a two-dimensional collective-variable space by a distance-based PCA. Within this PC1–PC2 plane, 50 representative seed conformations were selected to cover the landscape evenly rather than over-sampling the densest predicted region: a kernel density estimate discarded the sparsest predictions, and an inverse-density-weighted farthest-point-sampling scheme then chose seeds that maximized coverage while remaining anchored to the populated regions.

#### System setup and equilibration

The denist system was assembled with the CHARMM-GUI Membrane Builder: ^64,65^ the protein was embedded in a POPC lipid bilayer, solvated with TIP3P water, ^66^ and neutralized with ∼0.15 M NaCl. The CHARMM36m force field ^67^ was used for the protein, lipids, and ions. After energy minimization, the system was equilibrated using the standard six-step CHARMM-GUI protocol, in which positional and dihedral restraints on the protein and lipid headgroups were progressively released.

#### Relay target MD

The seeds were ordered along a minimum-spanning-tree relay path in PC space so that each seed could be reached from a structurally adjacent neighbor. Starting from the equilibrated highest-density structure, short restrained (target) MD runs (1 ns per segment) drove the system to each successive seed conformation using a Colvars ^68^ root-mean-square-deviation harmonic restraint (force constant 5 × 10^4^ kJ mol^−1^ nm^−2^, target center 0). The converged coordinates of each segment (typically within ∼1 Å of the targets) served as the starting structures for unbiased production MD.

#### Production MD

Production simulations were run in GROMACS 2025.4^69^ with a 4 fs time step enabled by hydrogen-mass repartitioning, ^70^ constraining all bonds to hydrogen with LINCS. ^71^ The temperature was held at 310.15 K with the velocity-rescaling thermostat ^72^ (τ_*T*_ = 1 ps) and the pressure at 1 bar using the C-rescale barostat ^73^ with semi-isotropic coupling (τ_*P*_ = 5 ps). Electrostatics were treated with particle-mesh Ewald ^74^ (1.2 nm real-space cutoff) and van der Waals interactions with a force-switched cutoff between 1.0 and 1.2 nm under a Verlet neighbor scheme. Each of the 50 seeds was simulated for 50 ns, resulting in 2.5 μs of aggregate sampling per system.

#### Free-energy surface and basin extraction

All production frames were projected onto the frozen PCA eigenvectors derived from the prediction ensemble. A two-dimensional Gaussian kernel density estimate of the projected population was converted to a free-energy surface via Δ*G* = −*k*_B_*T* ln ρ, and local minima of the surface were identified as conformational basins. For each basin, the production frame closest to the basin center in PC space was extracted as its representative structure for downstream structural comparison with experimental references.

## Data and code availability

The source code and data for SteerAF, reference-free state selection, PSSM analysis, and MD simulations will be available upon publication.

## Acknowledgements

The authors thank Ruihan Dong for suggesting the name *SteerAF*. This work was supported by the National Key R&D Program of China (2024YFA0916800) and the Innovative Research Group Project of the National Natural Science Foundation of China (T2321001). Part of the model inference and MD simulations was performed on the computing platform of the Center for Life Sciences at Peking University.

## Author Contributions

Conceptualization: Z.Z., C.S.; Methodology: J.T., Z.Z., S.Y., C.S.; Software: J.T., S.Y.; Validation: Z.Z.; Formal Analysis: J.T., S.Y.; Investigation: J.T., Z.Z., S.Y.; Resources: C.S.; Data Curation: J.T.; Writing – Original Draft: J.T., Z.Z., S.Y., C.S.; Writing – Review & Editing: J.T., Z.Z., S.Y., C.S.; Visualization: J.T., Z.Z., S.Y.; Supervision: C.S.; Project Administration: Z.Z., C.S.; Funding Acquisition: C.S.

## Supplementary Information

### 1 Supplementary Tables

**Table S1:** Hyperparameters of SteerAF.

| Symbol | Description | Default |
| --- | --- | --- |
| $T$ | Number of optimization steps | 6 |
| $N$ | Number of runs | 10—20 |
| $K$ | Warm-up steps (no recycling) | 3 |
| $\text{thr}_{\text{mask}}$ | Distogram mask threshold | 4 |
| $K_p$ | Step at which pLDDT loss is activated | 3 |
| $\lambda_p$ | pLDDT loss weight | 0.50 |
| $\rho$ | BGA sample ratio | 5% |
| lr | Learning rate (SGD step size) | $2 \times 10^{-2}$ |

**Table S2:** Curated PGK1 residue sites used to evaluate the biological relevance of SteerAF MSA-feature modifications.

| Site | Validated perturbation | Functional region | Role in open-closed conformational selection |
| --- | --- | --- | --- |
| R38 | R38A; R38K<sup>1,2</sup> | 3PG-binding side, N-domain catalytic site, substrate-signal input | Core domain-closure trigger. Mutational, kinetic, DSC, and SAXS evidence indicate that R38 is essential for induced domain closure. |
| K219 | K219A<sup>1</sup> | Nucleotide-binding side, nucleotide-site communication residue | Communication site linking the nucleotide-binding site to the main molecular hinge. |
| N336 | N336A<sup>1</sup> | ADP-side allosteric communication pathway | Coupling site between nucleotide binding and domain closure or closed-conformation stabilization. |
| E343 | E343A<sup>1</sup> | Nucleotide-site communication residue | Supports signal transmission from the nucleotide-binding site to the main hinge. |
| F165 | F165A<sup>3</sup> | Interdomain hinge-supporting region | Maintains hinge-region conformational integrity rather than acting as an isolated closure trigger. |
| E192 | E192A<sup>3,4</sup> | Main hinge and allosteric pathway hinge point | Key conformational-coupling residue in the main hinge region. |
| F196 | F196A<sup>3</sup> | Region near beta-strand L | Auxiliary regulator of domain movement within the hinge-neighbor mutant set. |
| S392 | S392A; S392A/T393A<sup>3</sup> | Beta-strand L and L14 hinge region | Regulates main-hinge geometry and substrate-coupled closure together with T393. |
| T393 | T393A; S392A/T393A; T393del<sup>3</sup> | Beta-strand L and L14 hinge region | Strong conformation-sensitive hinge site whose local geometry affects substrate-induced domain closure. |
| D218 | D218A; D218A/D374A<sup>5</sup> | MgATP/MgADP-binding side, phosphate-chain mobility regulator | Modulates open-state nucleotide binding and pre-closure flexibility through Mg<sup>2+</sup> and phosphate-chain interactions. |
| D374 | D374A; D218A/D374A; D374N; D374K<sup>5,6</sup> | MgATP/MgADP-binding side, ligand cooperativity, closed or TSA-like state regulator | Regulates the open-closed energy landscape and formation of fully closed or transition-state-analogue-like (TSA-like) complexes. |
| K215 | K215A; K215R<sup>1,2,6</sup> | Catalytic lysine, transferring-phosphate positioning, closed active-site geometry | Closed-state catalytic-geometry site; important for phosphate positioning but not a direct domain-closure trigger. |

**Table S3:** Curated hβ_2_AR residue sites used to evaluate the biological relevance of SteerAF MSA-feature modifications.

| Site | Validated perturbation | Functional region | Role in active-state conformational selection |
| --- | --- | --- | --- |
| D130 / D3.49 | D130N, D130A; combined mutations with E268<sup>7</sup> | DRY motif; TM3 cytoplasmic end; ionic-lock network | D130N/D130A disrupt the inactive-state TM3-TM6 ionic constraint. D130N is also associated with increased C285 accessibility and TM6 outward movement. |
| E268 / E6.30 | E268Q, E268A; D130N/E268Q; D130N/E268A<sup>7</sup> | TM6 cytoplasmic end; TM3-TM6 ionic lock | Together with D130 and R131, E268 forms the inactive-state ionic lock; E268Q/E268A single mutants and D130/E268 double mutants disrupt this lock, shifting the conformational equilibrium toward the active-like state. |
| L272 / L6.34 | L266S/K267R/H269K/L272A; L266S/L272A<sup>8</sup> | ICL3-TM6 cytoplasmic junction | Identified as part of a constitutively active mutant (CAM) combination that induces ligand-independent receptor activation. |
| F282 / F6.44 | F282L, F282A, F282G<sup>9</sup> | PIF/connector motif; TM6 helical core | F282L, F282A, and F282G single-point mutations produce varying degrees of constitutive activity and generate structurally distinct activated receptor conformations. |
| C285 / C6.47 | C285T<sup>10</sup> | TM6 proline-kink / rotamer toggle-switch region | C285T produces a constitutively active receptor indicating that C285 influences active-state conformational selection through the TM6 proline-kink / rotamer toggle switch. |

### 2 Supplementary Figures

**Figure S1:**
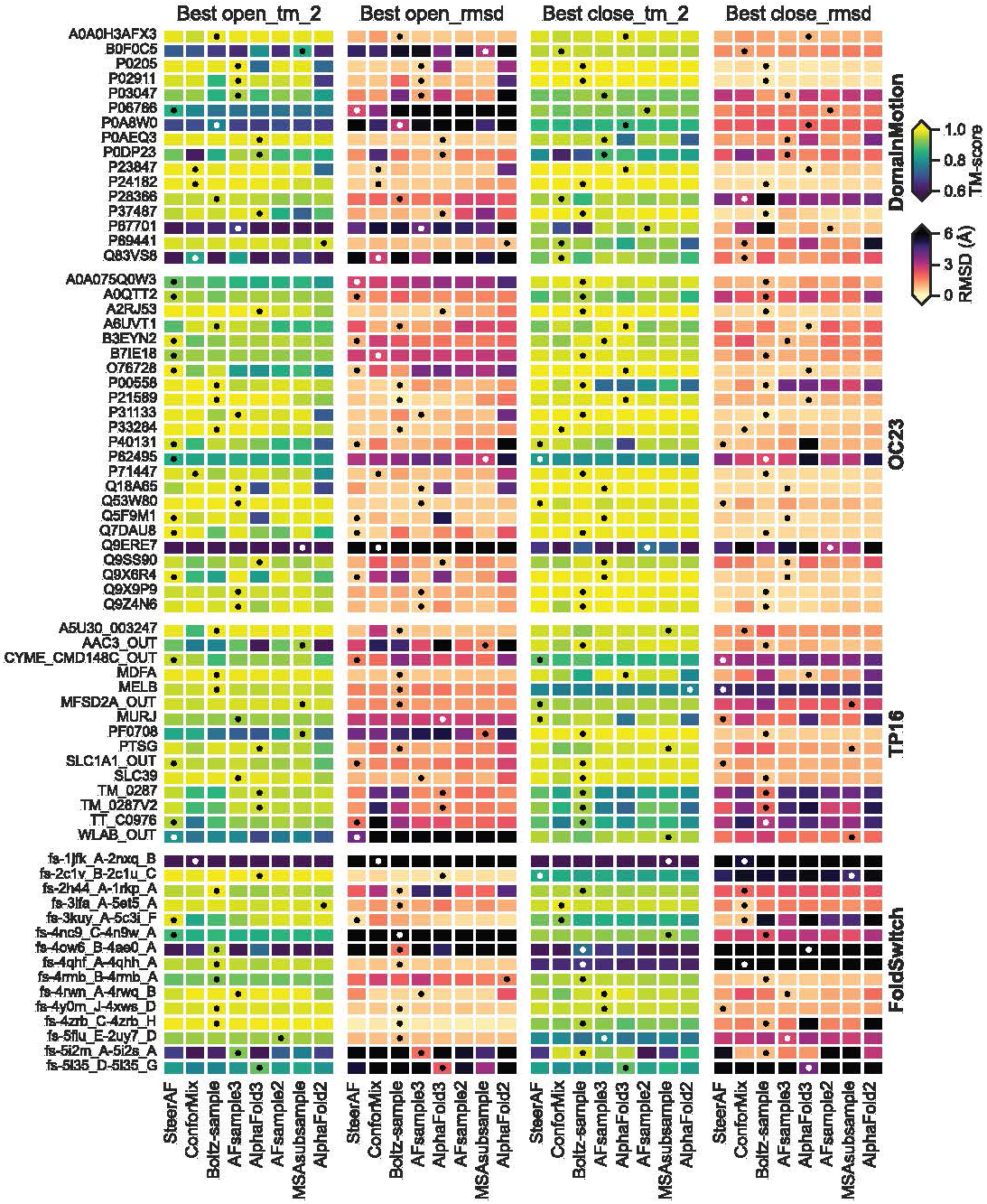
Per-system comparison of different methods. Black or white circles indicate the best method for each conformation.

**Figure S2:**
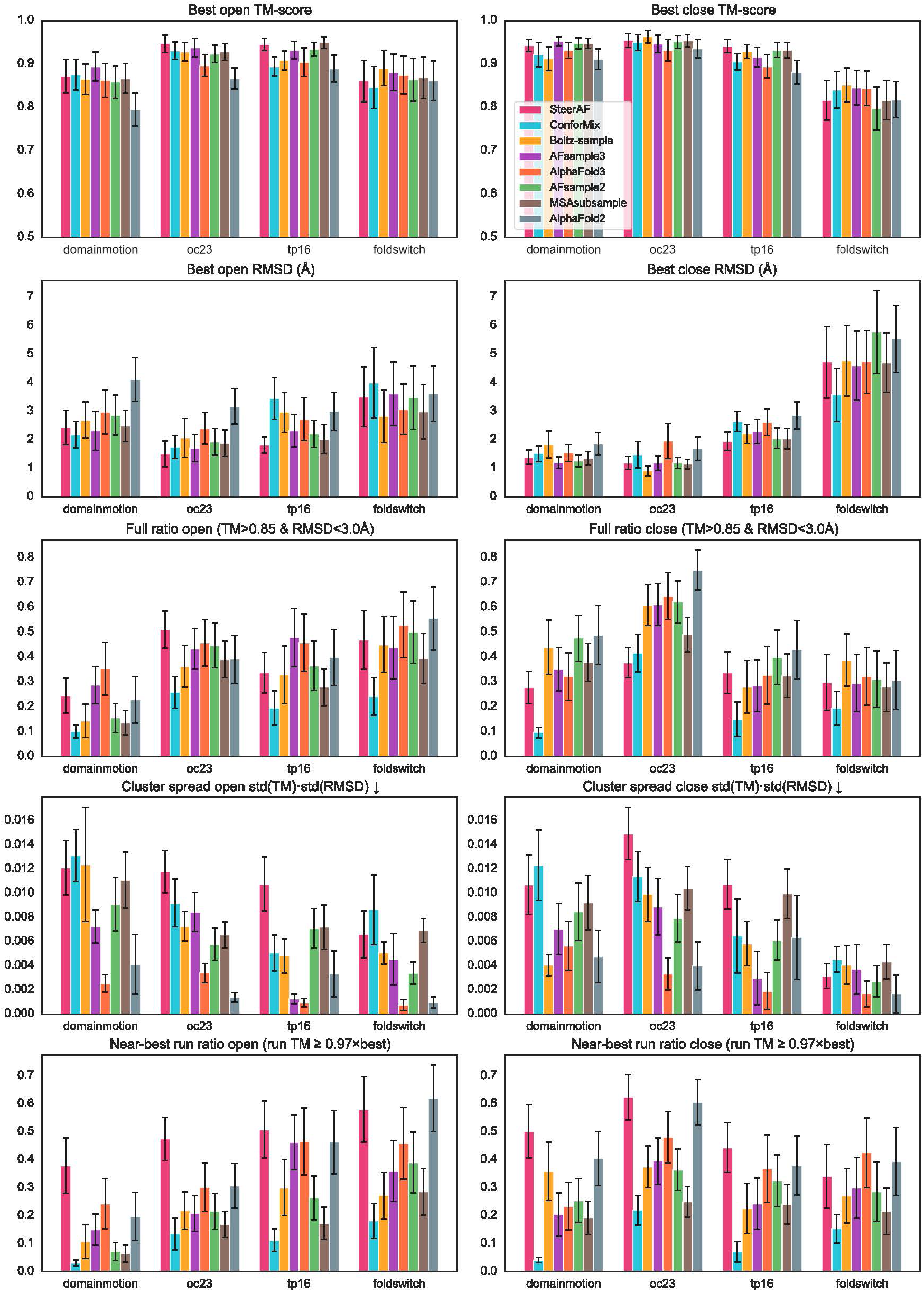
Metric comparison of different methods.

**Figure S3:**
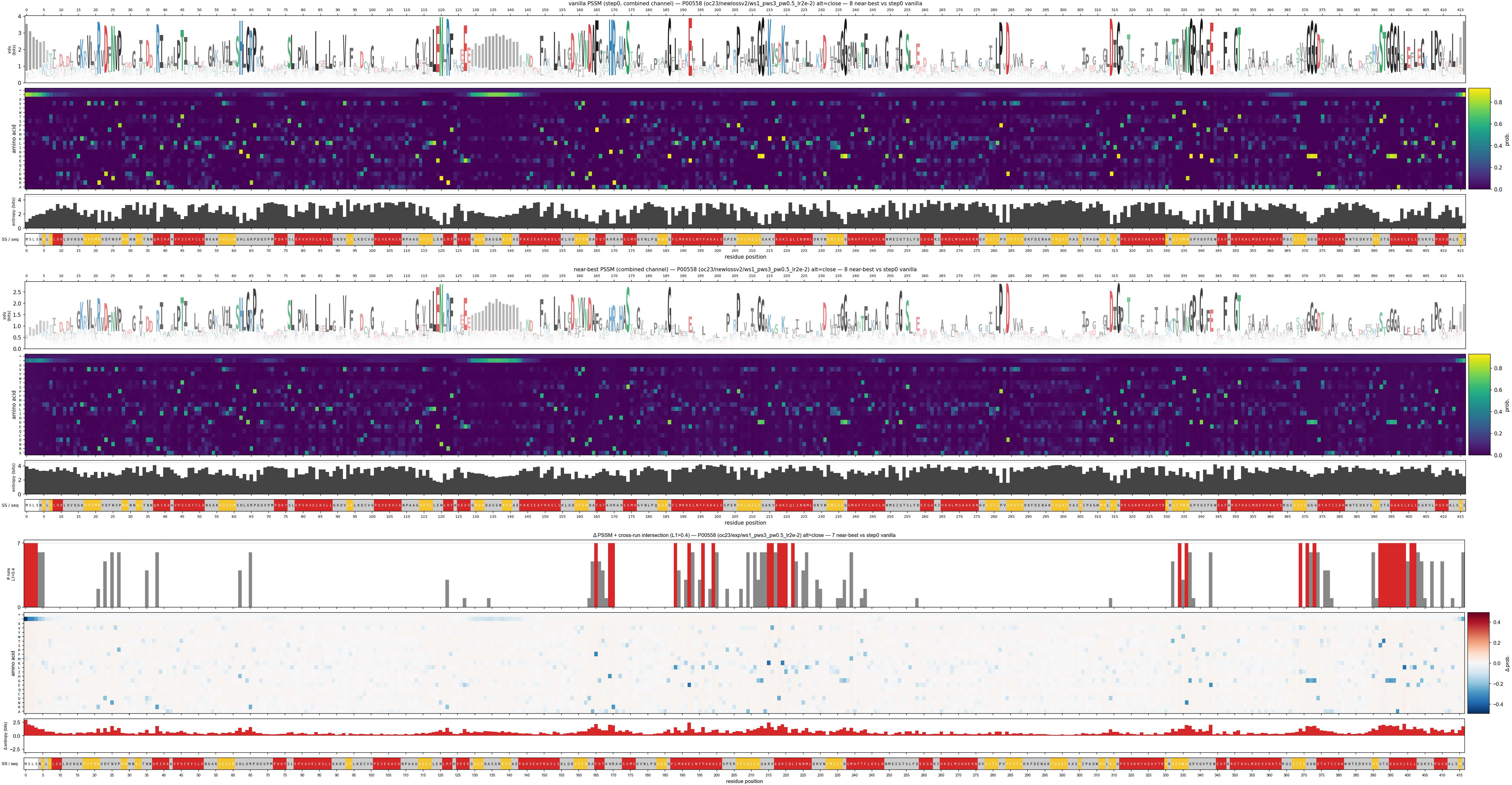
Stitched PSSM heatmaps for PGK1. The left, middle, and right blocks correspond, respectively, to the PSSM of the MSA feature at step-0, the PSSM of the MSA feature at the last near-best step (averaged over all near-best runs), and the difference between these two PSSMs (i.e., the PSSM of the MSA-feature modification). Within the left and middle blocks, the tracks from left to right are: the per-position sequence logo, ^11^ the PSSM heatmap, the per-position entropy of the PSSM, and a bar showing the secondary structure together with the query sequence. In the sequence logo, the total stack height at each position is the information content (in bits) at that position; in the entropy track, the value is the Shannon entropy of the amino-acid distribution at that position. The secondary structure is taken from the experimental structure of the default conformation: red denotes helices, yellow β-sheets, gray loops, and white positions unresolved in the experimental structure. The right (difference) block has no sequence logo; its tracks from left to right are a per-position count bar, the difference PSSM heatmap, the per-position entropy, and the same secondary-structure/query-sequence bar. The count bar gives, at each position, the number of the 7 last near-best predictions whose

**Figure S4:**
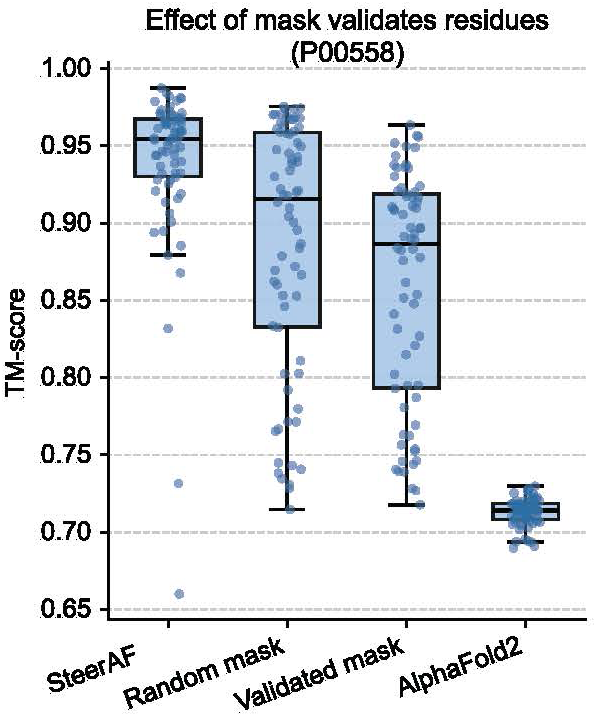
Site-restoration test on PGK1. For the PGK1 system, the MSA features of the literature-reported functional sites and of randomly selected control sites were each restored at the last near-best step to their step-0 values, and the TM-score distributions of the alternative-conformation predictions after restoration were compared between the two groups.

**Figure S5:**
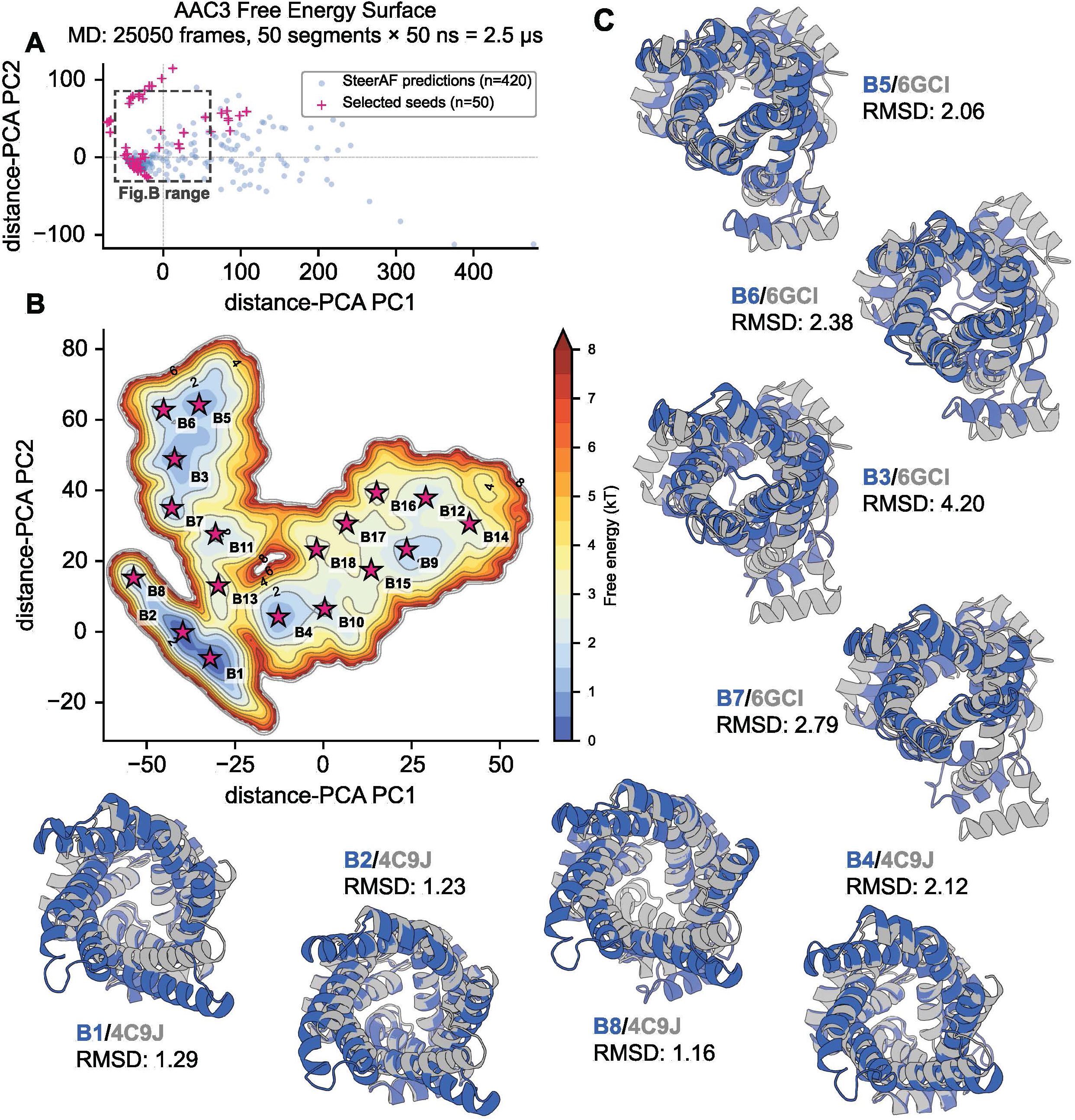
SteerAF predictions as starting points for MD-based free-energy exploration on AAC3. The same MD protocol used for MdfA was applied to the mitochondrial ADP/ATP carrier AAC3, recovering a multi-basin free-energy landscape that broadly expands the sampled conformational space relative to the prediction set. However, not all free-energy minima correspond to native folds. The low-energy basins reproduce the experimentally determined carrier states—the cytoplasmic-open c-state (PDB 4C9J; basins B1, B2, B8) and the matrix-open m-state (PDB 6GCI; basins B5, B6, B7)—to within ∼1–3 Å Cα RMSD. By contrast, several of the higher-energy, peripheral basins arise from partially or incorrectly folded structures, characterized by locally unwound or splayed transmembrane helices and large deviations (>4 Å) from any experimental reference. These misfolded minima are a generic by-product of seeding MD from an aggressively diversified prediction set, which by design includes low-confidence, non-physical conformations; on the relatively short simulation timescale they remain kinetically trapped and therefore appear as spurious basins. They are nonetheless readily identified—and excluded from functional interpretation—by their elevated free energy, peripheral location and abnormal pLDDT distribution in the distance-PCA space (Figure 3A), underscoring that the seeded-MD landscape should be read together with these structural-quality filters rather than treating every basin as a physical state.

**Figure S6:**
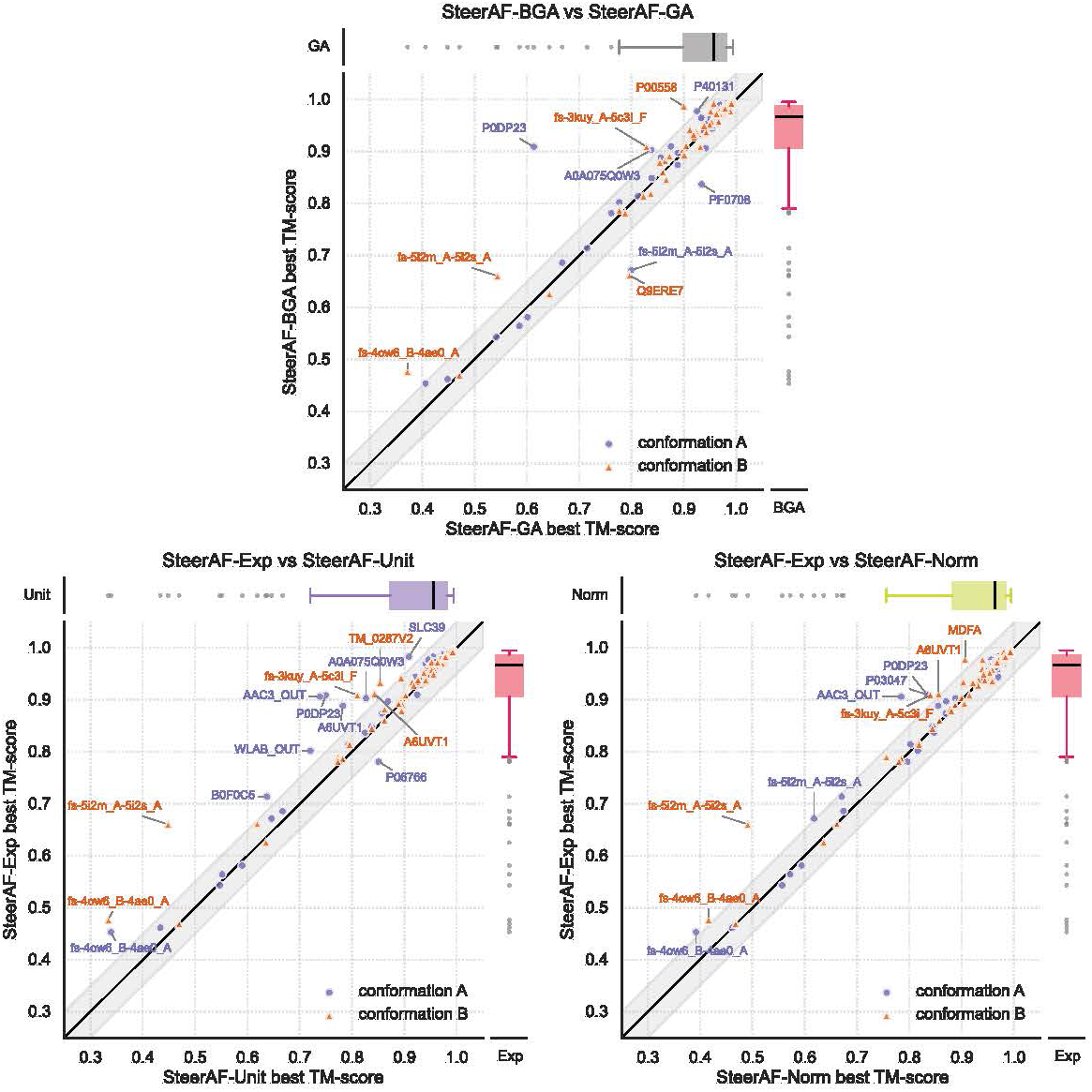
Comparison of different gradient ascent types and three gradient-regularization schemes

**Figure S7:**
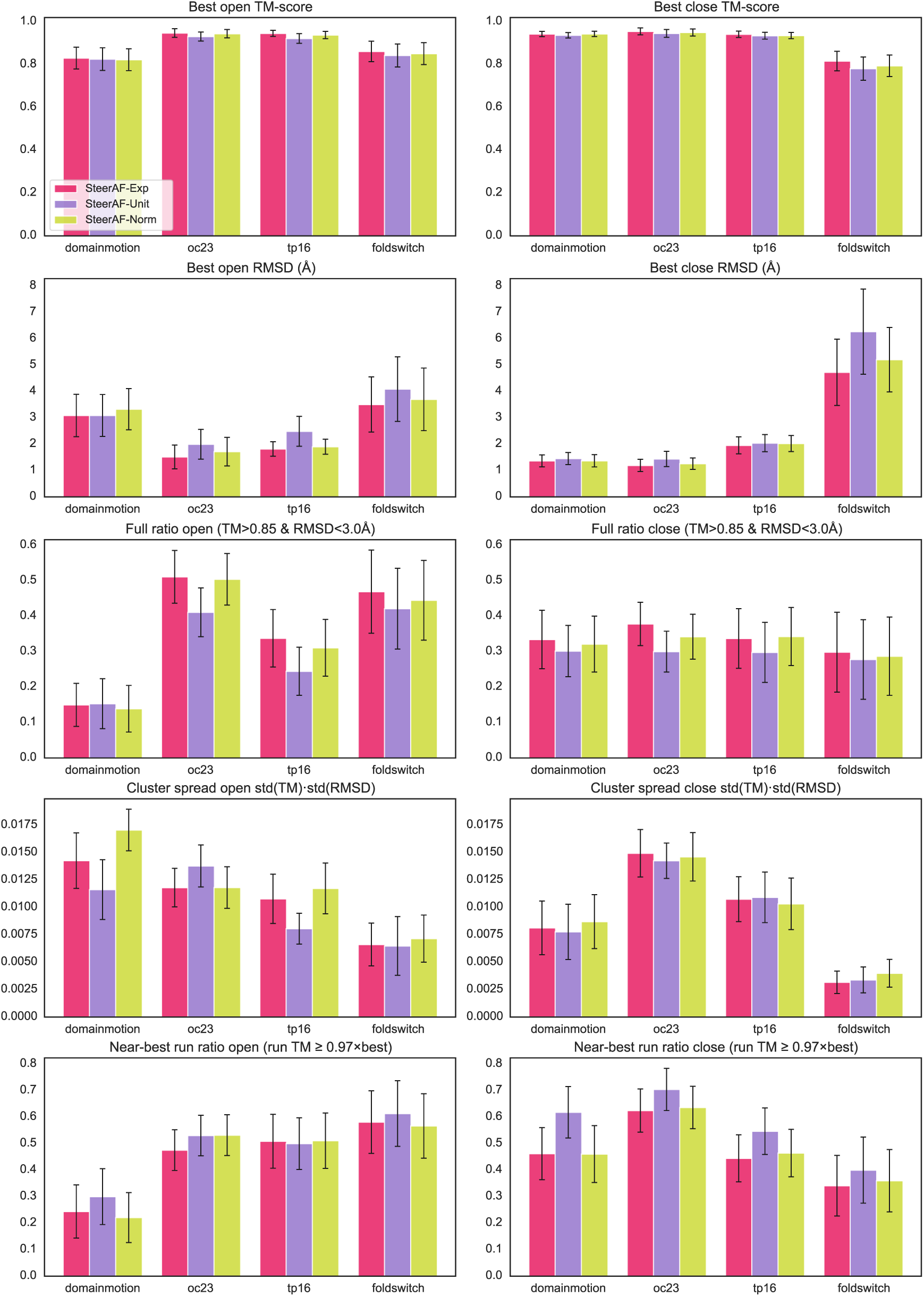
Metric comparison of the three gradient-regularization schemes

**Figure S8:**
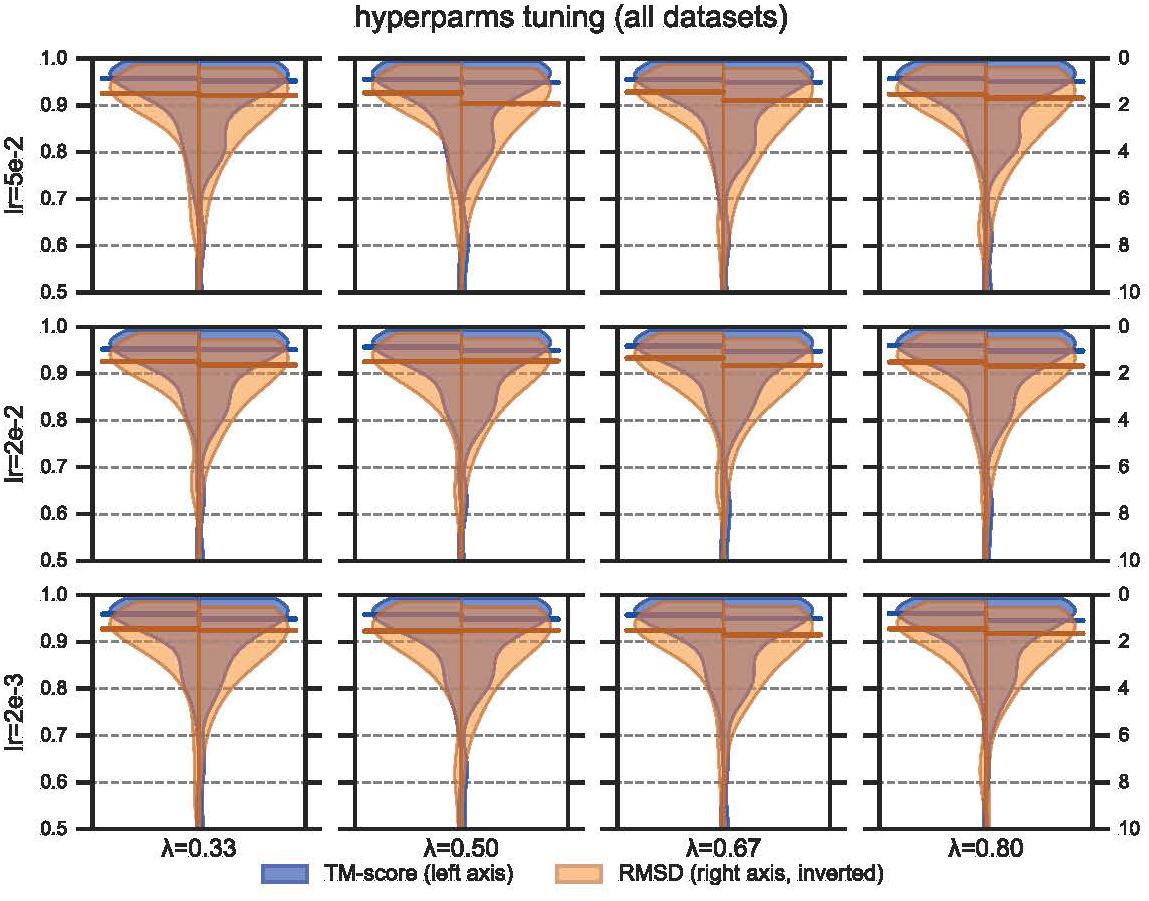
learning-rate and pLDDT-weight hyperparameter tuning.

